# Multi-scale coordination of planar cell polarity in planarians

**DOI:** 10.1101/324822

**Authors:** Hanh Thi-Kim Vu, Sarah Mansour, Michael Kücken, Corinna Blasse, Cyril Basquin, Juliette Azimzadeh, Eugene Wimberly Myers, Lutz Brusch, Jochen Christian Rink

**Affiliations:** Max Planck Institute for Molecular Cell Biology and Genetics, Pfotenhauerstrasse 108, 01307 Dresden, Germany.; Technische Universität Dresden, Zentrum für Informationsdienste und Hochleistungsrechnen (ZIH), Helmholtzstrasse 10, 01069 Dresden, Germany.; Institut Jacques Monod, Bâtiment Buffon, 15 rue Hélène Brion, 75205 Paris CEDEX 13, France.; Center for Systems Biology Dresden, Pfotenhauerstrasse 108, 01307 Dresden, Germany.

**Keywords:** Planaria, planar cell polarity, ciliary rootlet polarization, Fat/Dachsous, core PCP

## Abstract

Polarity is a universal design principle of biological systems that manifests at all organizational scales. Although well understood at the cellular level, the mechanisms that coordinate polarity at the tissue or organismal scale remain poorly understood. Here, we make use of the extreme body plan plasticity of planarian flatworms to probe the multi-scale coordination of polarity. Quantitative analysis of ciliary rootlet orientation in the epidermis reveals a global polarization field with head and tail as independent mediators of anteroposterior (A/P) polarization and the body margin influencing mediolateral (M/L) polarization. Mathematical modeling demonstrates that superposition of separate A/P- and M/L-fields can explain the global polarity field and we identify the core planar cell polarity (PCP) and Ft/Ds pathways as their specific mediators. Overall, our study establishes a mechanistic framework for the multi-scale coordination of planar polarity in planarians and establishes the core PCP and Ft/Ds pathways as evolutionarily conserved 2D-polarization module.

## Introduction

Polarity is a fundamental design principle of biological systems. The end-to-end assembly of structurally asymmetric tubulin dimers generates micron-length microtubules with end to end polarity, the directional movement of motor proteins and their cargo along polarized microtubules contributes to the structural polarization of cells, and neighbor-neighbor coupling of the polarity vectors of individual cells generates tissue-level polarization. Finally, in the case of *Bilaterians*, the positioning of the brain and eyes in front and the tail at the rear signifies polarity also at the level of the organismal body plan. Yet the invariant head-tail alignment of scales in the fish skin or polarity patterns of wing hairs in the fruit fly wing demonstrates the coordination of polarity vectors between the organismal and ultimately molecular levels. How this is achieved remains a fascinating problem.

Polarity is best understood at the cell and tissue level, owing to the genetic analyses of wing hair patterns in *Drosophila* (Aw and Devenport, 2017; Lawrence et al., 2007; Yang and Mlodzik, 2015) and cilia orientation in vertebrates (Wallingford, 2010). Both scenarios involve the activity of the evolutionarily conserved core PCP pathway (Adler, 2012; Butler and Wallingford, 2017; Hale and Strutt, 2015; Yang and Mlodzik, 2015), which encompasses transmembrane proteins such as Frizzled (Fz), VanGogh (Vang) and the atypical cadherin Flamingo (Fmi), intracellular proteins Disheveled (Dvl), Diego (Dgo) and Prickle (Pk) in addition to multiple accessory effector components. The core PCP pathway generates polarity by the mutual antagonism between two sub-complexes, the distal Fz/Dvl/Dgo complex and the proximal Vang/Pk complex. Within individual cells, they establish mutually exclusive Fz/Dvl or Vang/Pk membrane domains. At the tissue level, the preferential interaction between Fz/Fmi dimers of one cell and the Vang/Fmi dimers of its neighbor aligns polarization vectors between cells and results in the emergence of tissue-scale PCP-polarity patterns. A parallel pathway involving the atypical cadherins Fat (Ft) and Dachsous (Ds) and the intracellular components Dachs and Four-jointed (Fj) establishes cell and tissue polarity in a conceptually similar manner and the two pathways interact to varying degrees during fly wing development (Lawrence et al., 2007; Matis and Axelrod, 2013). Finally, functional coupling between subcellular polarity landmarks and actin polymerization or ciliary rootlet orientation translate into the consistent alignment of wing-hairs in *Drosophila* or the head-tail alignment of cilia in the *Xenopus* larval skin (Wallingford, 2010). With individual cells as autonomously polarizing units and neighbor-neighbor coupling as propagation mechanism, the tissue-wide coordination of polarization vectors and their invariant alignment with the cardinal body axes constitute ongoing conceptual and mechanistic challenges in the field.

Simulations of the PCP machinery suggest an inherent tendency of neighbor-neighbor coupled systems to get trapped in “local energy minima” swirl patterns (Amonlirdviman, 2005; Burak and Shraiman, 2009; Hazelwood and Hancock, 2013) and long-range cues such as morphogen or gene expression gradients have been proposed as mechanism for ensuring tissue-scale polarity coordination (Strutt, 2009; Yang and Mlodzik, 2015). In the *Drosophila* fly wing, core PCP and Ft/Ds polarity vectors first orient towards the anteroposterior (A/P) and dorsoventral (D/V) organizers and the emergence of the final wing hair polarity pattern involves dynamic vector reorientations between these landmarks at different stages of development (Sagner et al., 2012). The D/V and A/P organizers also provide sources of patterning signals and ectopic sources of the D/V patterning signal Wingless (Wg) can re-orient polarity (Wu et al., 2013). Further, core components of the Ft/Ds pathway are expressed in gradients along the pathway’s main axis of polarization (Ma et al., 2003; Zeidler et al., 2000). However, the rescue of Ft mutant polarity phenotypes by uniformly overexpressed transgenes (Matakatsu, 2004; Matakatsu and Blair, 2006; 2012; Matis et al., 2014) and likely low or no diffusion of the highly lipophilic Wnt ligand (Alexandre et al., 2014) both challenge the significance of gradients as long-range polarization mechanism. Mechanical shear stress associated with tissue morphogenesis has been shown to orient PCP vectors in the *Drosophila* wing (Aigouy et al., 2010) and the *Xenopus* epidermis (Chien et al., 2015). However, core PCP components in turn may affect the shear-causing cell flows during vertebrate gastrulation (Butler and Wallingford, 2017), which makes the cause-consequence relationship between PCP-orientation and mechanical stress difficult to disentangle. Finally, mathematical modeling indicates that long-range cues might not be required for long range polarization in systems that grow from a small size (Aigouy et al., 2010). Hence neither the origins nor long-range propagation of polarity patterns are well understood at present.

The multi-scale coordination of polarity is mainly studied “bottom up”, i.e., by alterations at the molecular level and examination of tissue scale consequences. “Top-down” approaches such as the experimental reversal of the organismal A/P axis and examination of the ensuing tissue and cell level consequences are hardly possible with the current developmental model systems. In search of a new system allowing such multi-scale probing of polarity, we here make use of the striking body plan plasticity of planarian flatworms. Adult planarians can be induced to convert their tail into a head, to sprout ectopic heads all along the body margins or to regenerate a tail instead of a head (Gurley et al., 2008; Iglesias et al., 2008; Petersen and Reddien, 2008). Such anatomical plasticity is rooted in the continuous renewal of all cell types via constantly dividing pluripotent adult stem cells (neoblasts) and the requisite need for patterning signals to mediate location-dependent cell fate choices (Rink, 2012). A Wnt signaling gradient emanating from the tail is a core component of planarian A/P patterning (Reuter et al., 2015; Stückemann et al., 2017; Sureda-Gómez et al., 2016) and experimental interference with Wnt signaling consequently result in the spectacular reprogramming of organismal head/tail polarity. Additionally, the cilia-driven directional gliding motility of planarians provides clear indications of cell-level polarity. The ventral epidermis of planarians is largely comprised of multiciliated cells and the worms locomote via the concerted beating of cilia (Glazer et al., 2010; Rink et al., 2009; Rompolas et al., 2013). Cilia-driven motility in turn requires the alignment of individual cilia within and between all cells of the epithelium, which can be visualized by the orientation of the evolutionarily conserved macromolecular rootlet that anchors cilia to the cytoskeleton (Frisch and Farbman, 1968; Park et al., 2008; Yang et al., 2002). With ciliary rootlet orientation as generic polarity readout and global body plan polarity as an experimental variable, planarians therefore offer a unique experimental system to examine the multiscale coordination of polarity.

We have developed the requisite antibody, imaging and image analysis tools and find that planarians indeed display global polarization of cilia orientation along the head-tail axis, albeit with a lateral splay component in the head region. By reprogramming body plan polarity at the organismal level, we find that the head tip, tail tip and the body edge provide autonomous polarization cues. We use mathematical modeling to demonstrate their collective sufficiency for explaining the topology of the epidermal polarity field as the propagation of boundary conditions. Further, we identify the core PCP pathway as specific mediator of the A/P polarization component and the Ft/Ds pathway as specific mediator of the mediolateral (M/L) polarization component. Overall, our study establishes a mechanistic framework for the multi-scale polarity coordination in planarians and the core PCP and Fat/Ds as evolutionarily conserved 2D-polarization module.

## Results

### Visualization of ciliary rootlet polarity

The ventral epidermis of planarians is comprised of regularly spaced multi-ciliated cells, with their protruding axonemes forming a dense lawn of cilia (Fig. 1A). While polarity cannot be inferred from the random orientation of the long and slender axonemes (Fig. 1A), work in other model systems has established in several instances that the macromolecular rootlet that anchors each cilium in the cell cortex is aligned with the direction of the ciliary beat (Frisch and Farbman, 1968; Park et al., 2008; Rieger, 1981; Yang et al., 2002). To ascertain that this is also the case in planarians, we first carried out thin section electron microscopy of the ventral epidermis. Ciliary rootlets were readily apparent and displayed the stereotypical striated rootlet extension and a basal foot projecting in the opposite direction from the basal body (Fig. 1B). Importantly, grazing section further revealed parallel alignment among individual rootlets within and between individual epidermal cells (Fig. 1C), thus establishing the principal utility of ciliary rootlet orientation as polarity readout in planarians.

**Figure 1.**
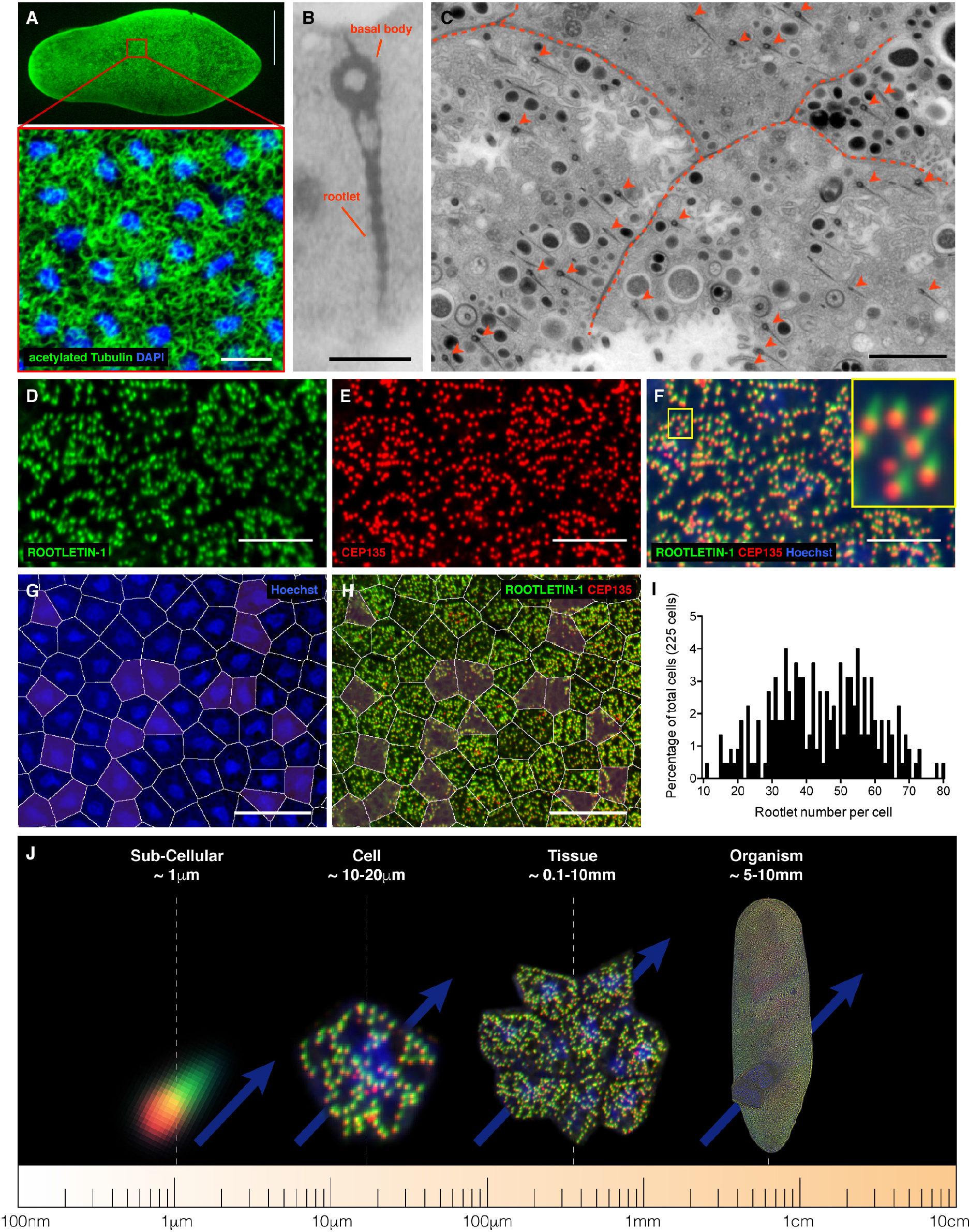
Ciliary rootlets as indicators of planarian tissue polarity. **A.** Whole mount acetylated tubulin immunostaining showing the axonemes (green) of the densely ciliated ventral epidermis. Scale bar: 500 μm. Zoomed view: additional nuclear staining (DAPI, blue), illustrating the regular spacing of epithelial cells. Scale bar: 10 μm. **B.** TEM image of a single ciliary rootlet, sectioned along the long axis. Scale bar: 500 nm. **C.** TEM image of a grazing section of the ventral epidermis. Dashed lines: Cell-cell boundaries. Arrowheads: Individual ciliary rootlets. Scale bar: 2.2 μm. **D-F.** Rootlet visualization by confocal microscopy, using antibody staining against SMED-ROOTLETIN-1 (D) and SMED-CEP135 (E), overlaid with nuclear staining (Hoechst) in the merged image (F; color coding as before). Magnified inset: Visualization of rootlet orientation by chevron-shaped ROOTLETIN signal and the dot-like basal body staining at the broad end. Scale bar: 10 μm. **G.** Computational cell boundary approximation by Voronoi tessellation (white outlines) of nuclear staining (DAPI, blue). **H.** ROOTLETIN-1 (green) and CEP135 (red) antibody staining of the same cell field as in G, illustrating generally good agreement between actual and inferred cell outlines with the exception of non-ciliated small cells (purple overlay). Their equal tessellation tends to underestimates the size of neighboring cells. Images are single confocal sections. Scale bar: 25 μm. **I.** Quantification of cellular rootlet number, based on manual counting of 225 cells from 6 wild-type animals. Cell boundaries were inferred by Voronoi tessellation as in G. **J.** Illustration of multi-scale polarity analysis afforded by our combined imaging and processing pipeline. See text for details.

Faced with the need to visualize the orientation of micron-length rootlets across centimeter-long planarians, we turned to light microscopy. We made use of an existing antibody against the basal body component SMED-CEP135 (Azimzadeh et al., 2012) and additionally raised a monoclonal antibody against SMED-ROOTLETIN-1, one of the two planarian homologues of the highly conserved structural Rootletin protein (Fig. S1A-B) (Yang et al., 2002). Both antibodies label abundant punctate structures in ventral epidermal cells (Fig. 1D-E), which resolve into dot-like CEP135-positive centrosomes situated at the broad end of chevron-shaped ROOTLETIN-1-positive structures (Fig. 1F). The staining patterns of the two markers therefore faithfully recapitulate the rootlet architecture observed by electron microscopy (Fig. 1B) and further allow the visualization of rootlet orientation by light microscopy, as previously demonstrated in *Xenopus* larval skin (Park et al., 2008). The planarian epidermis consists mainly of regularly spaced multiciliated cells ~ 10-15 μm in diameter (Fig. 1G-H), which harbor ~ 40 rootlets on average (Fig. 1I). Previous BrdU pulse/chase experiments demonstrated that the planarian epidermis undergoes continuous cell turnover (Tu et al., 2015; van Wolfswinkel et al., 2014). The interspersed nonciliated cells (Fig. 1H) might therefore represent integrating epidermal progenitors, which suggests that the planar polarization of the planarian epidermis is a dynamic process, similar to the vertebrate skin (Devenport et al., 2011). As pre-condition for investigating polarity coordination across the tissue, we developed an imaging pipeline comprising tiled Z-stack volume acquisition on a fast spinning disk confocal microscope, automated stitching of the resulting image stacks, epidermal layer extraction, rootlet detection and directionality quantification (Fig. S1C; (Blasse et al., 2017)). Verification with manually annotated ground truth data confirmed both sensitivity and accuracy of our rootlet analysis pipeline (Fig. S1D-H). Overall, these efforts provide a toolkit for the multi-scale analysis of polarity in planarians (Fig. 1J).

### Polarity field in wild type planarians

At the organismal level, the polarization of the planarian body plan is evident on the basis of the position of the eyes in the head and distinct marker gene expression domains in the head, trunk and tail (Fig. 2A). At the tissue and cell levels, we investigated polarity by using our pipeline to visualize rootlet orientation across the entire ventral epidermis (Fig. 2B). Averaging rootlet orientation within 100 *μ*m squares (~ 80 cells/3000 rootlets) and plotting the resulting direction vector for each square revealed consistent long-range polarization along the A/P axis, consistent with the cilia driven gliding motility of planarians (Fig. 2C). Further, especially in the head region, tissue polarity vectors deviated towards the body margins (referred to as “splay” in the following text). Rootlet orientation at the sub-cellular level is therefore clearly coordinated with the planarian body plan polarity at the organismal level. Thus, rootlet orientation provides an effective readout of cell and tissue polarity, much like the hairs in the *Drosophila* wing (Adler, 2012) or the ciliary rootlets in the *Xenopus* larval skin (Park et al., 2008).

**Figure 2.**
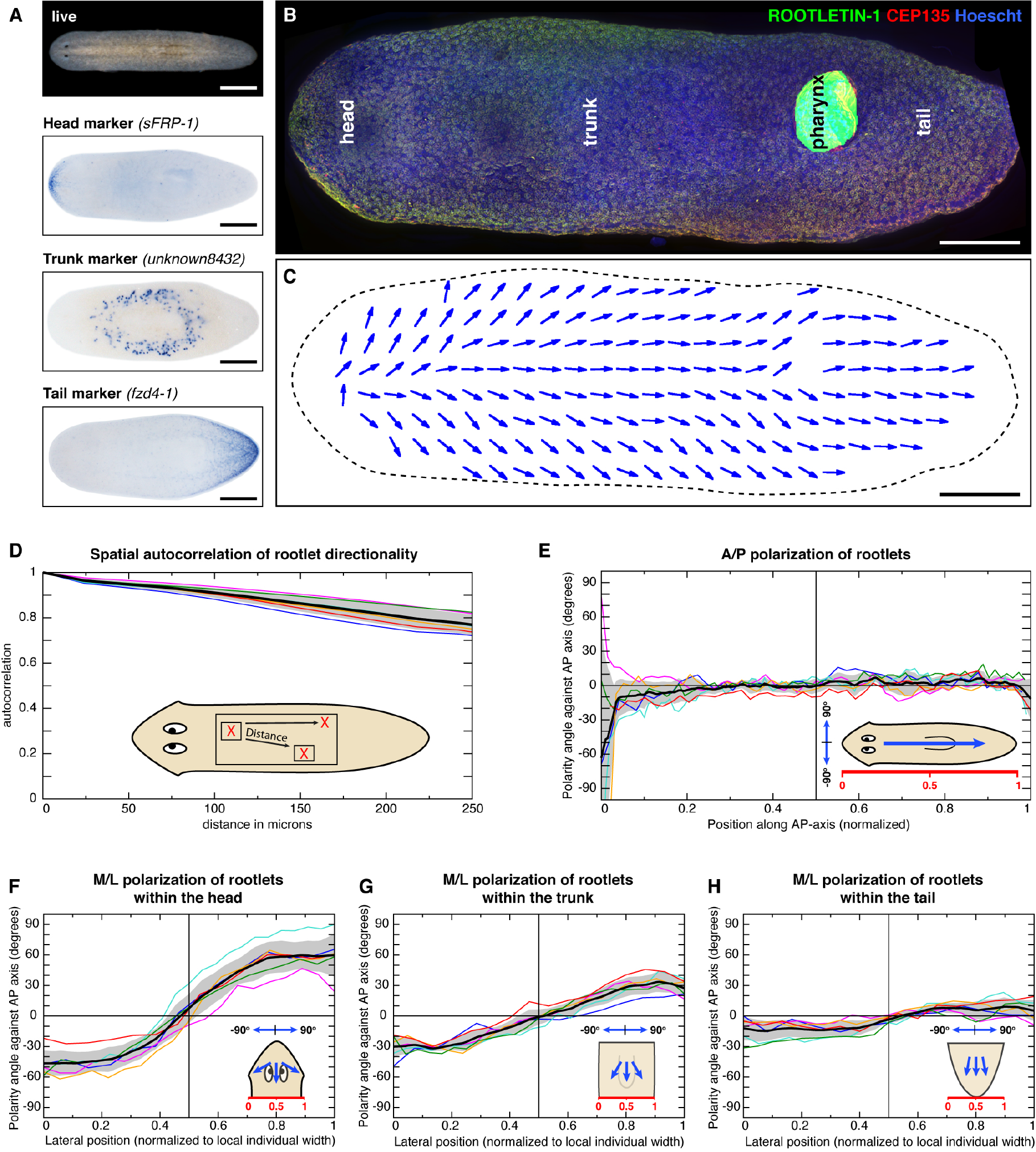
Rootlet orientation reveals the organismal polarity field of the ventral epidermis. **A.** Body plan polarity landmarks in wild type *S. mediterranea*. Live image (top), and whole mount in situ hybridization gene expression patterns of head, trunk and tail markers (bottom). Scale bar: 500 μm. **B.** Stitched and surface extracted high-resolution image of the entire ventral epidermis as used for rootlet orientation quantification. Anti-ROOTLETIN-1 (green), anti-CEP135 (red) and nuclear staining (Hoechst, blue). Scale bar: 250 μm. **C.** Vector field representation of the average ciliary rootlet orientation within 100 μm squares of the specimen shown in B. Arrow direction corresponds to the centrosome-rootlet vector. Scale bar: 250 μm. **D.** Spatial autocorrelation analysis of rootlet directionality along the A/P axis, comparing directional coherence within squares of 150×150 pixels as a function of distance (μm). **E.** A/P polarization component: Averaged rootlet orientation orthogonal to the midline along the A/P axis in 6 wild type animals, traces were normalized by length. **F-H.** M/L polarization component: Averaged rootlet orientation parallel to the midline along the M/L axis in the head (F), trunk (G) and tail (H) of 6 wild type animals. Traces were normalized by width. D-H: Colored lines represent individual specimen. The solid black line and the grey shading indicate mean plus/minus standard deviation of 6 animals, respectively. 0° designates head to tail orientation, +/− 90° orthogonal orientations and cartoons illustrate measured rootlet orientations.

To quantitatively assess this polarization of the ventral epidermis, we took advantage of the very large number of individual rootlet quantifications (~1-2 millions rootlets/animal) and processing capability (typically 6 animals/condition) that our pipeline affords. We first assessed the coherence of local rootlet orientation as a function of distance. The measured spatial autocorrelation values of average cilia orientation within 150×150 pixels windows (~16 cells) remained above 80% within 200 μm (Fig. 2D), which quantitatively confirms the neighbor-neighbor coupling of cellular polarization vectors (Fig. 1C) and the long-range coordination of cell polarity suggested by the coarse-grained view (Fig. 2C). To analyze local rootlet orientation in relation to global body plan polarity, we defined 0° as signifying perfect alignment with the head/tail axis, 180° as opposite tail/head orientation and 90° or 270° as orthogonal orientation to the midline. Averaging rootlet orientation within bins orthogonal to the midline quantitatively confirmed the consistent and highly reproducible orientation of cilia along the head/tail axis, with the deviations at the head and tail tips resulting from folding artifacts of the whole mount preparations (Fig. 2E). Averaging rootlet orientation within bins parallel to the midline quantitatively confirmed the above noted splay (Fig. 2F-H). In the head (Fig. 2F), rootlets at the body margins partially orient towards the body edge (~ −45° left margin, 45° right margin), while rootlets in the vicinity of the planarian midline are perfectly oriented along the head/tail axis. The highly reproducible sigmoidal shape of the M/L orientation traces further revealed that the magnitude of the splay progressive decreases along the A/P axis, averaging +/− 30° in the trunk region (Fig. 2G) and decreasing to <10° in the tail region (Fig. 2H). Overall, our analysis reveals a globally coordinated polarity field comprised of components aligning with the cardinal A/P and M/L axes of the planarian body plan.

### Global coordination of local cell polarity

To identify the global cues that determine local cell and tissue polarity, we next made use of the unique body plan plasticity of planarians. RNAi of *Smed-β-catenin-1* in regenerating trunk pieces forces regeneration of an ectopic head instead of a tail (Gurley et al., 2008; Iglesias et al., 2008; Petersen and Reddien, 2008) and thus results in double-headed animals with a complete reversal of organismal A/P polarity in the posterior body half (Fig. 3A). Intriguingly, double-headed *β-catenin-1(RNAi)* animals displayed bipolar polarity fields (Fig. 3B, S2A-C) with a 180° reversal of cell and tissue polarity approximately at the midpoint of the body (Fig. 3C). This further implies that even cells in the pre-existing tissue (pigmented area in Fig. 3A) underwent a complete polarity reversal. These results therefore demonstrate that the planarian head is the source of a long-range polarity cue that orients cell polarity away from the body margin.

**Figure 3.**
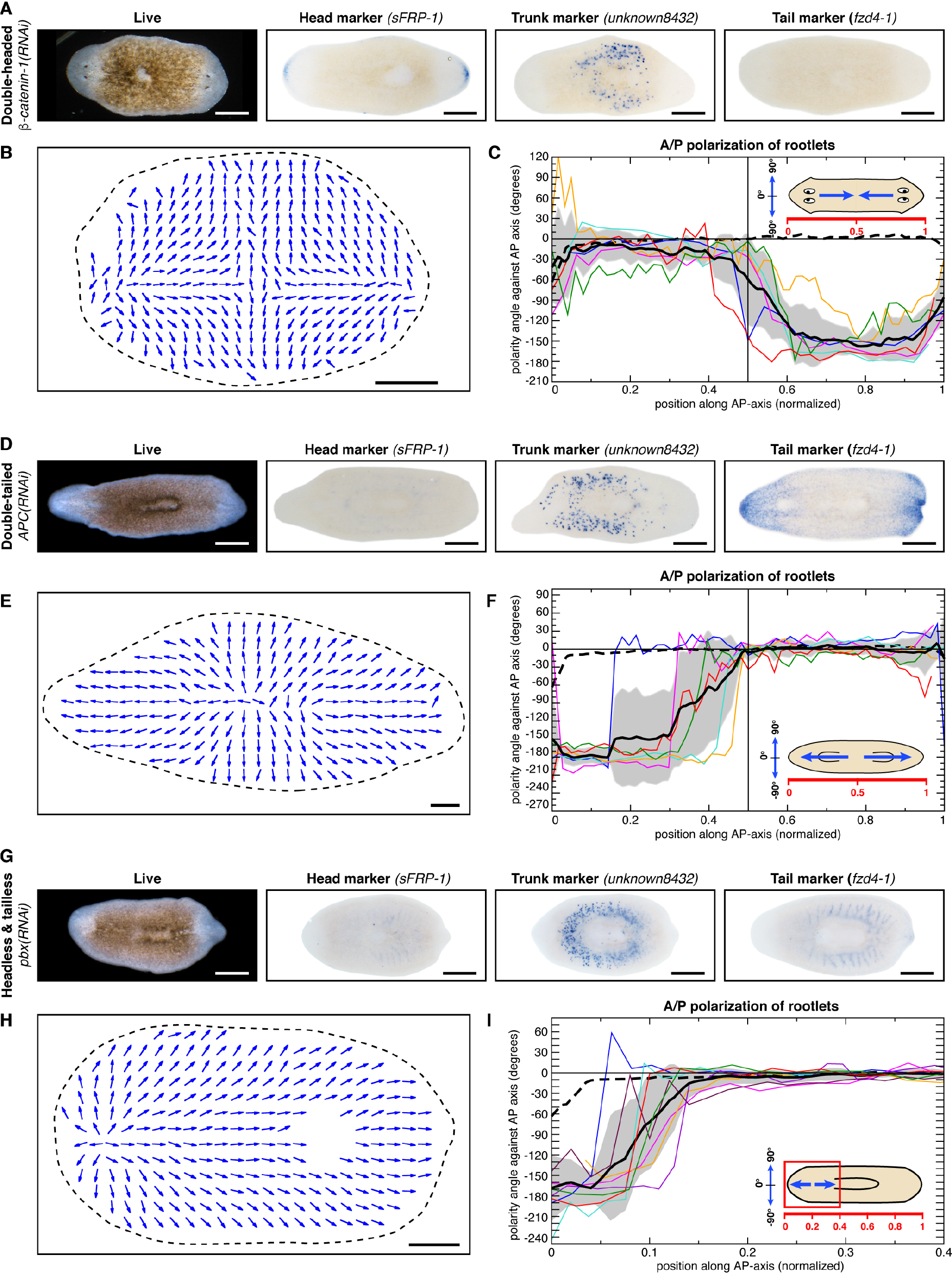
Head, tail and the body margin provide polarity cues. **A.** Double-headed animals resulting from the bipolar regeneration of *β-catenin-1(RNAi)* trunk pieces. Left: Live image. Right: Head, trunk, and tail marker expression visualized by whole mount in situ hybridization. Scale bar: 500 μm. **B.** Vector field representation of the average ciliary rootlet orientation within 100 μm bins of a representative double-headed specimen. Scale bar: 250 μm. **C.** A/P-polarization component in double-headed specimens: Lines trace the averaged rootlet orientation orthogonal to the midline along the A/P axis in 6 double-headed animals, traces were normalized by length. **D.** Double-tailed animals resulting from the bi-polar regeneration of *APC(RNAi)* trunk pieces. Left: Live image. Right: Head, trunk, and tail marker expression visualized by whole mount in situ hybridization. Scale bar: 500 μm. **E.** Vector field representation of the average ciliary rootlet orientation within 100 μm bins of a representative double-tailed specimen. Scale bar: 250 μm. **F.** A/P-polarization component in double-tailed specimens: Lines trace the averaged rootlet orientation orthogonal to the midline in 6 double-tailed animals, traces were normalized by length. **G.** Headless and tailless animals resulting from the regeneration of trunk pieces under *pbx(RNAi)*. Left: Live image. Right: Head, trunk, and tail marker expression by whole mount in situ hybridization. Scale bar: 500 μm. **H.** Vector field representation of the average ciliary rootlet orientation within 100 μm bins of a representative double-headed specimen. Scale bar: 250 μm. **I.** A/P-polarization component in headless and tailless specimens: Lines trace the averaged rootlet orientation orthogonal to the midline in 6 headless and tailless animals, traces were normalized by length. C, F, I: Colored lines represent individual length-normalized specimens. The solid black line and grey shading indicate mean plus/minus standard deviation, respectively. The dashed line represents the average of 6 wild type specimens. 0° designates head to tail orientation, +/− 90° orthogonal orientations and cartoons illustrate measured rootlet orientations.

To ask whether the planarian tail might also provide a polarity cue, we made use of the double-tailed animals resulting from elevated canonical Wnt signaling under *Smed-APC(RNAi)* (Fig. 3D). Indeed, double-tailed planarians displayed a bipolar polarity field (Fig. 3E, S2D-F) with an invariant 180° polarity reversal along the A/P axis (Fig. 3F). Intriguingly, however, polarity vectors were always oriented towards the body margins and thus in the opposite direction of the double-headed animals. This result demonstrates that the planarian tail also provides a long-range polarity cue and that the head-tail orientation of the wild type polarity field therefore likely originates from the synergistic interplay between the two global organizers.

To assess polarity in the absence of both A/P polarity cues, we made use of the headless and tailless animals resulting from the knockdown of *Smed-pbx* (Fig. 3G) (Blassberg et al., 2013; Chen et al., 2013). *pbx(RNAi)* prevents formation of the head and tail poles that are thought to act as organizers of the planarian A/P axis (Oderberg et al., 2017; Reuter et al., 2015; Vásquez-Doorman and Petersen, 2014; Vogg et al., 2014). Remarkably, *pbx(RNAi)* animals still displayed coherent long-range polarity (Fig. 3H) with a high degree of local order (Fig. S3B). Neither the predominant A/P-orientation (Fig. 3G) nor the lateral splay (Fig. S3C) was affected throughout most of the field, suggesting the maintenance of pre-existing tissue polarity even in the absence of global cues. However, *pbx(RNAi)* regenerates invariably displayed a specific reversal of tissue polarity at the “anterior” end (Fig. 3I, Fig. S3D). This finding was intriguing, as it indicated the possibility of a further “attractive” orienting effect of the body margin on rootlet orientation, which would consequently antagonize pre-existing tissue polarity at the anterior, but not the posterior end of regenerated trunk pieces.

Overall, our manipulations of organismal body plan polarity establish head, tail and possibly also the body margin as organizers of the epidermal polarity field.

### Model development, parametrization and validation

To explore whether these cues are sufficient for explaining the global polarity field, we turned to mathematical modeling. Although our model is inspired by the importance of compartment boundaries in fly wing polarity establishment (Sagner et al., 2012), we deliberately take a coarse-grained approach that does not make any assumptions regarding the underlying molecular mechanisms. Rather, our model is built on three fundamental assumptions (Fig. 4A). First, the local fixation of polarity vectors by head, tail and body edge acts as boundary conditions. Second, the boundary vectors propagate via neighbor-neighbor coupling into the interior of the cell field. Third, the orientation of ciliary rootlet is a faithful readout of the combined effect of all polarity cues. Specifically, the polarity field 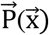 as the model’s actual representation of rootlet orientation emerges from the superposition of two independent polarity components, the A/P field 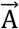 governed by the synergistic effects of the head and tail boundary vectors, and the radially directed M/L field 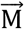 governed by the body margin. Both fields are defined on a two-dimensional domain approximating the planarian body shape, which renders the precise value of 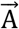 and 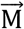 a function of the spatial position 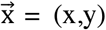. Mathematically, 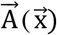 and 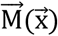 are defined by partial differential equations with no flux boundary conditions and boundary-restricted reaction terms (Fig. 4B and Supplemental materials), the superposition of which generates the field 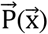. Mathematically more parsimonious models with only one polarity field can also qualitatively explain multiple features of the observed polarization patterns (see Supplemental materials), but are less well supported by experimental evidence (see below and discussion). Overall, our model allows the quantitative analysis of the interplay between local polarity and global cues, as well as the prediction of the dynamic consequences of experimental perturbations (Fig. S4A-H)

**Figure 4.**
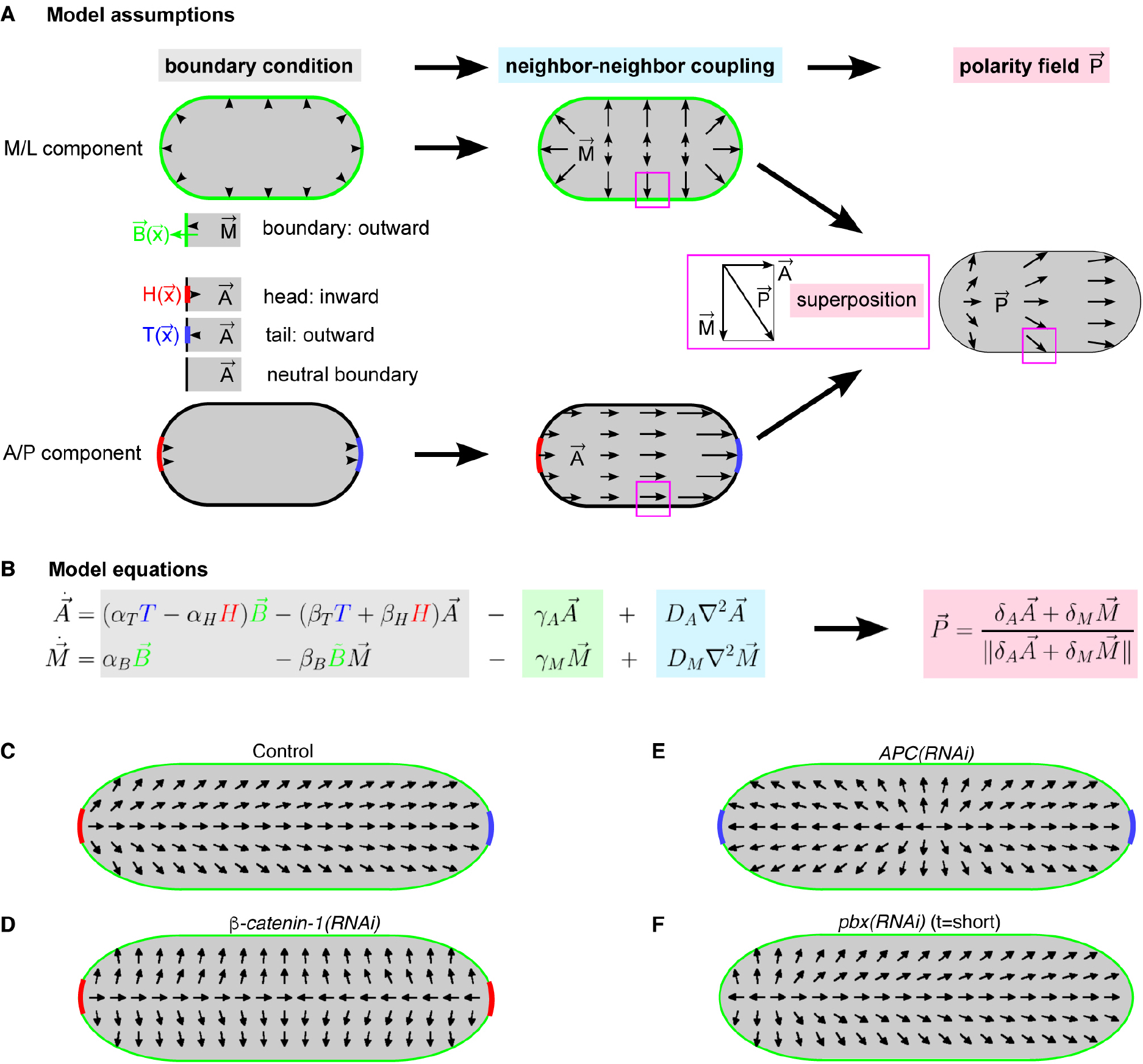
Boundary-cue model of ciliary rootlet polarization. **A.** Boundary conditions (left): The planarian body margin (top) is represented by the coordinate vector 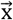 (green), which is modeled by the outward-pointing vector field 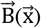 that has modulus 1 on the boundary and vanishes elsewhere. The localized influence of the head (red) and tail (blue) are defined by scalar fields 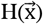 and 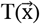 that are 1 at the indicated position of the boundary and 0 elsewhere. Neighbor-neighbor coupling (center), transmits the boundary cues to the cell field and generates the two separate polarity fields 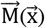 (radially directed) and 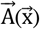 (A/P-directed). Linear superposition of 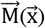 and 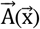 (pink, right section) finally results in the unit vector field 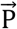 as the model’s representation of actual ciliary rootlet orientation. **B.** Model equations for the temporal dynamics of 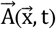, 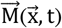 and 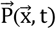. Two partial differential equations for 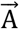 and 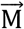 determine their temporal change (left hand side, dots denote time derivative) from three terms (colored boxes) that technically implement the above processes. The first term (gray) generates values for 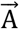 and 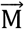, exclusively at boundary locations set by 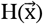, 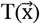 and 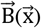. Changes in the ratios α_*H*_/*β*_*H*_, α_*T*_/*β*_*T*_ and α_*B*_/*β*_*B*_ change the magnitude of 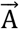 and 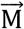, thus allowing the simulation of gradual and dynamic changes of boundary values in response to experimental perturbations. The opposite vector orientation of 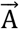 at head versus tail is represented by the respective signs of the α’s and the factor 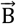. The operator B̃ coordinate-wise multiplies 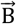 to 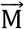. The second term (green) implements a spatially uniform depolarization with rate γ (e.g., due to dynamic cell turnover), which is a prerequisite for dynamic remodeling of existing polarity. The third term (blue) is the diffusion operator with coefficient D which implements the spatially averaging effect of neighbor-neighbor coupling. No-flux boundary conditions are used for this operator. Ciliary rootlet polarity 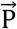 (pink box) is given by the weighted sum of 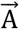 and 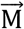 and normalization of nonzero values to modulus 1. Parameters δ_*A*_ and δ_*M*_ modulate the responsiveness of cells to the 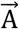 and/or 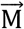 component. For model analysis and parameter estimation see Supplemental materials for detail explanation. **C-F.** Simulations of the experimentally observed wild type (C), double-headed (D), double-tailed (E) and headless and tailless (F) ciliary polarity fields. All simulations were initialized with the same wild type pattern and only the boundary conditions for the head and tail 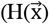, 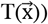 were changed. C-E represent final states, F a temporal intermediate of the simulation.

We first asked whether the model can qualitatively reproduce the polarity field of wild type planarians (Fig. 2C). Indeed, a combination of manual and data-based parameter inference (Fig. S4A) recapitulated both the predominant polarization of ciliary rootlets along the A/P axis, as well as the A/P graded lateral splay (Fig. 4C). The latter necessitates restriction of the head boundary effect (H) to the head tip, which corresponds to the presumed location of the planarian head organizer (Chen et al., 2013; Oderberg et al., 2017; Vásquez-Doorman and Petersen, 2014; Vogg et al., 2014) and thus has important implications regarding the alignment of polarity vectors with the cardinal body axes (see discussion). Further, recapitulation of the A/P gradation of the splay pattern despite the presumed globally uniform radial polarization effect of the body margin requires the tail to be more strongly polarizing than the head (α_*T*_/*β*_*T*_ > α_*H*_/*β*_*H*_), which then results in the graded relative contribution of the 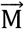 field along the A/P axis. The splay pattern thus emerges as secondary consequence of the primary assumptions of the model, which provides a first line of evidence for the validity of the underlying assumptions.

Next, we simulated the reversal of organismal polarity in double-headed animals. As shown in Fig. 4D, solely the substitution of the wild type tail pole (T) parameter settings by the corresponding head parameters (H) was sufficient for the recapitulation of the bi-polar polarity field *in β-catenin-1(RNAi)* animals (Fig. 3B). The substitution of the wild type head (H) parameters with the corresponding tail (T) settings recapitulated the opposite polarity reversal in double tailed *APC(RNAi)* animals, again without the need for any additional parameter tuning (Fig. 4E). Interestingly, the model also succeeded in qualitatively reproducing the localized polarity reversal at the anterior of “headless and tailless” *pbx(RNAi)* animals (Fig. 4F). Specifically, this effect emerges as a dynamic consequence of combined head and tail cue inactivation (α_*H*_, α_T_ = 0) and consequent local domination of the 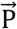 field by the now unopposed M/L polarity component. Therefore, this result provides additional evidence for the presumed polarity-organizing role of the body margin. In general, the close recapitulation of our experimental results by the model demonstrates that the combined boundary condition effects of head, tail and body margin are indeed sufficient for explaining the multi-scale coordination of polarity in the planarian epidermis.

### Ft/Ds mediates the M/L polarization component

A key prediction of our model is that the 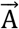 and 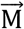 fields are functionally distinct. Therefore, separate molecular activities should exist that orient rootlets in A/P and M/L directions. Towards the goal of identifying the underlying mechanisms by loss of function screening, we first simulated the selective loss of the M/L polarization component (either by inactivation of the boundary cue, α_*M*_ = 0, or loss of the cellular response to it, δ_*M*_ = 0). This approach predicted the specific loss of the splay and consequently uniform A/P orientation of the polarity field as expected knockdown phenotype of M/L polarization components (Fig. 5A). We next asked whether the comparatively few molecular pathways that have thus far been shown to mediate planar cell polarization phenomena across the animal kingdom are present in *S. mediterranea*. Planarians harbor homologs of all core components of the Ft/Ds pathway, including the large protocadherins Dachsous (*Smed-ds*) and Fat (2 homologues, *Smed-ft-1* and *Smed-ft-2*), as well as the atypical myosin Dachs (two homologues, *Smed-dachs-1* and *Smed-dachs-2*) and the Golgi-localized kinase Four-jointed (*Smed-fj*). We found the genes to be broadly expressed in various cell types and organs, including prominent *ds*-expression along the body edge (Fig. 5B), and largely overlapping expression of different pathway members in the epidermis (Fig. S5A) (Wurtzel et al., 2015). Intriguingly, knockdown of the single *ds* homologue in intact animals resulted in a striking recapitulation of the M/L component loss prediction (Fig. 5C). Intact *ds(RNAi)* animals invariably displayed a reduced splay component all along the A/P axis (Fig. 5D), with corresponding statistically significant reductions in the slope of M/L polarity trajectories in head, trunk and tail (Fig. 5E). Interestingly, the decay of the spatial autocorrelation function and average rootlet polarization along the A/P axis were unaffected (Fig. S5C-D), indicating that Ds is neither required for rootlet orientation along the head/tail axis nor for the coordination of rootlet orientation *per se*. Together with the very similar phenotypes *of ft-1,-2(RNAi)* animals (Fig. S5E-F), these data strongly indicate that the planarian Ft/Ds pathway is specifically required for orienting ciliary rootlets relative to the body margins.

**Figure 5.**
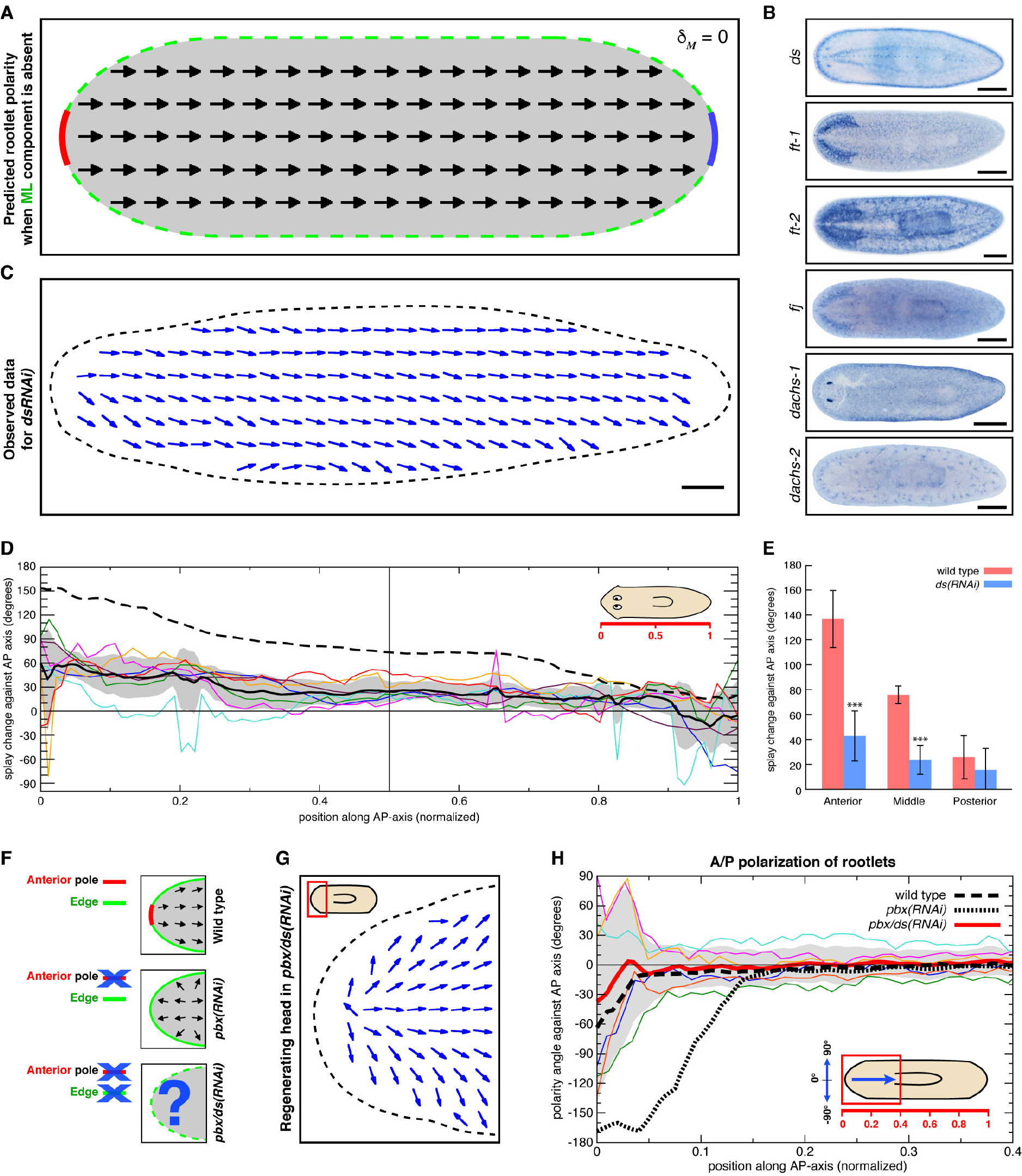
Ft/Ds is required for M/L component that orients ciliary polarity in planarians. **A.** Predicted vector field representation of average ciliary rootlet orientation under loss of the cellular response to the M/L component, (δ_*M*_= 0). **B.** Expression patterns of Ft/Ds pathway genes by whole mount in situ hybridization. Scale bar: 500 μm. **C.** Vector field representation of the average ciliary rootlet orientation within 100 μm bins of a representative *ds(RNAi)* specimen. Scale bar:
250 μm. **D.** Splay angle change along the A/P axis under *ds(RNAi)*, representing the slope of rootlet orientation change between left and right body margin as graphed in Fig. 2F-H. All traces were normalized by length. Colored lines represent 7 individual specimens. The solid black line and grey shading indicate the average plus/minus standard deviation, the dashed line the mean of 6 wild type specimens. 0° designates uniform A/P orientation, 180° orientation towards the margins. **E.** Quantitative comparison of splay angle changes between *ds(RNAi)* and wild type specimens, based on D. Splay angles of 7 specimens each were averaged within the head, trunk and tail territory. * p-value < 0.05; ** p-value < 0.01; *** p-value < 0.001. **F.** Rationale of the double-RNAi experiment to probe the organizing role of the body margin, see text for details. **G.** Vector field representation of average ciliary rootlet orientation within 100 μm squares in of a representative headless *pbx/ds(RNAi)* specimens. Cartoon: Displayed region. **H.** *ds(RNAi)* rescues the anterior specific polarity reversal induced by *pbx(RNAi)*. A/P polarization component in the anterior region of headless *pbx/ds(RNAi)* specimens. Colored lines trace the averaged rootlet orientation orthogonal to the midline along the A/P axis, traces were normalized by length. Thick, red line and grey shading indicate mean plus/minus standard deviation of 6 *pbx/ds(RNAi)* specimens, respectively. Thick dashed line represents the mean of 6 wild type specimens. Thick dotted line indicates mean of 6 *pbx(RNAi)* specimens. 0° designates head to tail orientation, +/− 180° the opposite directionality.

To test whether the Ft/Ds pathway indeed mediates the M/L polarity component in our model, we turned to “headless” *pbx(RNAi)* animals. Since we infer the unopposed dominance of the M/L component as cause of anterior polarity reversal, the concomitant knockdown of *ds* should rescue the effect (Fig. 5F). Strikingly, simultaneous RNAi of *pbx* and *ds* in regenerating animals indeed prevented the anterior polarity reversal observed after RNAi of *pbx* alone (Fig. 5G-H, Fig. S5G-I). However, *pbx* knockdown also restored the splay gradient along the A/P axis that *ds(RNAi)* alone abolishes (Fig. 5G, Fig. S5I). Simply combining the parameter settings of the single knock down simulations (reduced A/P component in *pbx(RNAi)* and reduced M/L component in *ds(RNAi)*; Fig. S4E) quantitatively reproduced this effect without any further parameter tuning. Overall, these results conclusively establish the role of the body margin as polarity organizer and the planarian Ft/Ds pathway as the mechanism that aligns ciliary rootlets accordingly. Further, they demonstrate that the splay effect depends on the relative ratio between M/L and A/P cues, rather than their absolute strength. As such, these data also provide experimental support for the superposition of A/P and M/L components as governing principle of rootlet orientation 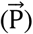 (Fig. 4A) and thus another key premise of our model.

### Core PCP signaling mediates A/P polarization

Towards the goal of also identifying molecular mediators of the A/P polarization component, we again simulated the expected RNAi phenotype of its total loss (either by removing the head and tail cues, α_*H*_, α_*T*_ = 0, or the ability of cells to respond to them, δ_*A*_ = 0). The resulting unopposed dominance of the M/L polarization component then results in the radial orientation of ciliary rootlets towards the closest boundary (Fig. 6A). In search of corresponding loss-of-function phenotypes, we also examined the conservation of the core PCP pathway in planarians as further highly conserved polarity pathway (Hale and Strutt, 2015). Planarians harbor homologues of many PCP core pathway members (e.g., a total of 9 Fz homologues (Liu et al., 2013; Stückemann et al., 2017), 3 Vang-homologues (this study and (Almuedo-Castillo et al., 2011)) and 2 Dvl-homologues (Gurley et al., 2008)) that are expressed in many cell types, including the epidermis (Fig. S6A). However, our search so far failed to identify clear homologues of the core pathway members Fmi/Celsr or Pk (not shown). We first focused our functional analysis on double knockdown of the two Dvl homologues due to Dvl’s absolute requirement for PCP and to minimize redundancy concerns (e.g., amongst the many Fz receptors). Ventral epidermal cells of *Smed-dvl-1,-2(RNAi)* animals displayed normal rootlet numbers (Fig. S6B) and also rootlet morphology appeared normal at the light microscopic level (Fig. 6B). This result was unexpected, given that in *Xenopus* Dvl is required for docking the ciliary rootlet (basal body) to the actin cortex (Park et al., 2008) and that a previous report had suggested similar roles for the planarian homologues (Almuedo-Castillo et al., 2011). However, *dvl-1,-2(RNAi)* animals lost their tail under our experimental conditions (Fig. 6C), which is consistent with the expected inhibition of the tail-maintaining canonical Wnt signal upon Dvl depletion (Gurley et al., 2008). *dvl-1,-2(RNAi)* animals also displayed a global radial re-polarization of rootlet orientation (Fig. 6D) in striking resemblance to the predicted consequence of A/P polarization component loss (Fig. 6A). The orientation of rootlets towards the body edge was strongly and significantly increased all along the A/P axis (Fig. 6E-F). Although the more rapid decay of the spatial autocorrelation amongst rootlets indicated weakened neighbor-neighbour coupling, *dvl-1,-2(RNAi)* animals clearly maintained coordinated rootlet polarization within and between epidermal cells (Fig. S6C). Overall, these phenotypes indicate a key role of planarian Dvl in the A/P orientation of ciliary rootlets all along the A/P axis.

**Figure 6.**
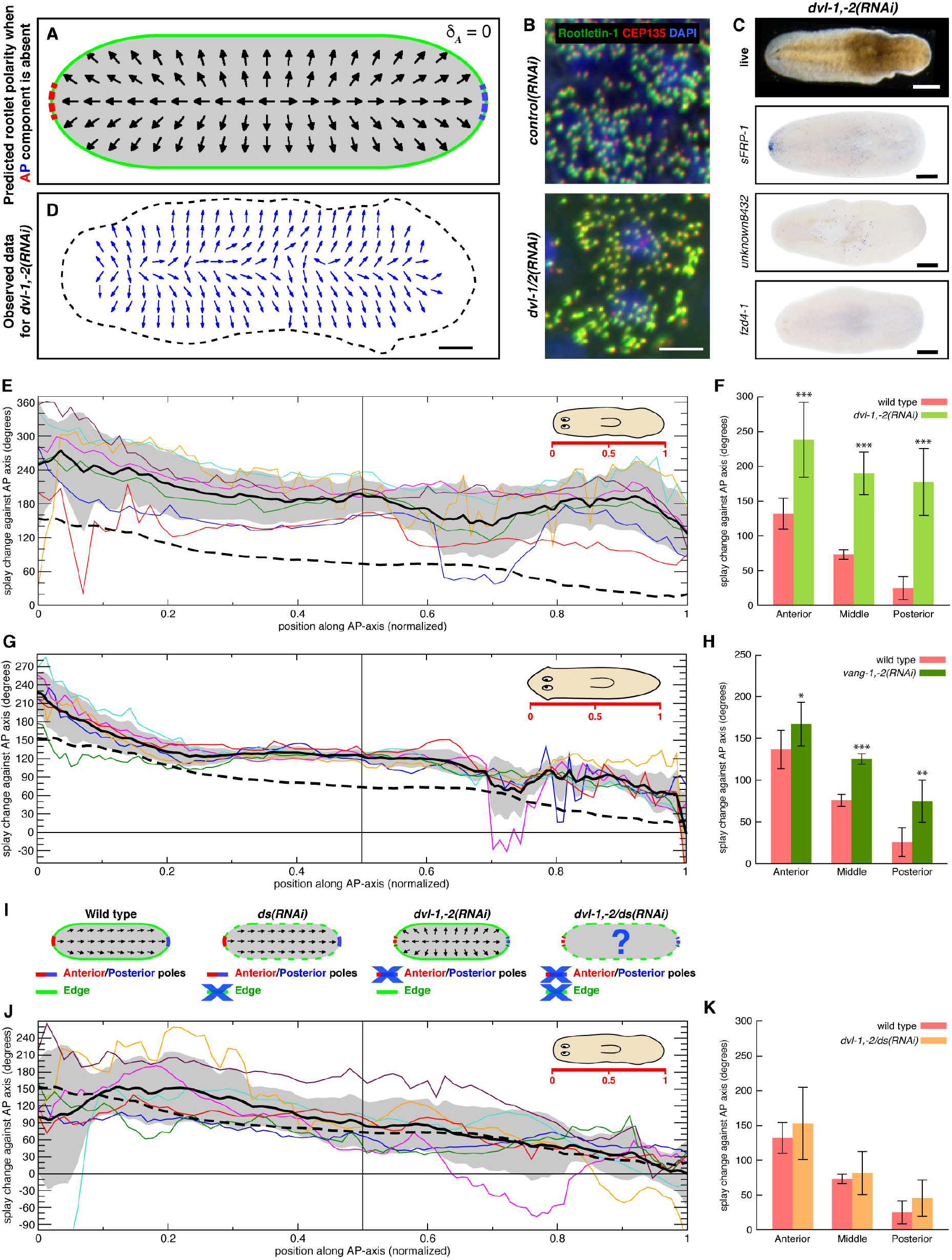
Core PCP is required for A/P component that orients ciliary polarity in planarians. **A.** Predicted vector field representation of average ciliary rootlet orientation under loss of the cellular response to the A/P polarity component (δ_A_ = 0). **B.** Ciliary rootlets in control and *dvl-1,-2(RNAi)*. ROOTLETIN-1 (green) and CEP135 (red) antibody staining; nuclei labeled with Hoechst (blue). Scale bar: 5 μm. **C.** Live-image of *dvl-1,-2(RNAi)* specimen (top), head, trunk, and tail marker expression by whole mount in situ hybridization below. Scale bar: 500 μm. **D.** Vector field representation of the average ciliary rootlet orientation within 100 μm squares of a representative *dvl-1,-2(RNAi)* specimen. Scale bar: 250 μm. **E.** Splay angle change along the A/P axis under *dvl-1,-2(RNAi)* as compared to wild type. **F.** Quantitative comparison of splay angle changes between wild type and *dvl-1,-2(RNAi)*, based on E. Splay angles of 7 specimens each were averaged within the head, trunk and tail territory. **G.** Splay angle change along the A/P axis under *vang-1,-2(RNAi)* as compared to wild type. **H.** Quantitative comparison of splay angle changes between wild type and *vang-1,-2(RNAi)*, based on G. Splay angles of 6 specimens each were averaged within the head, trunk and tail territory. **I.** Rationale of *dvl-1,-2/ds* triple-RNAi experiment, see text for details. **J.** Splay angle change along the A/P axis under *dvl-1,-2/ds(RNAi)* as compared to wild type. **K.** Quantitative comparison of splay angle changes between *dvl-1,-2/ds(RNAi)* and wild type, based on J. Splay angles of 7 specimens each were averaged within the head, trunk and tail territory. F, H, K: * p-value < 0.05; ** p-value < 0.01; *** p-value < 0.001. E, G, J: Colored lines represent individual specimens. The solid black line and grey shading indicate the mean plus/minus standard deviation, respectively. The dashed line represents the mean of 6 wild type specimens. 0° designates uniform A/P orientation, 180° M/L orientation.

Since Dvl is also an essential component of the canonical Wnt signaling pathway that patterns the planarian A/P axis (Gurley et al., 2008; Stückemann et al., 2017), the RNAi phenotype could reflect either a non-cell autonomous consequence of the observed tail cue loss (Fig. 6C) and/or a cell autonomous role in the transmission of planar cell polarization within and between epidermal cells. Simulations of the *dvl-1,-2(RNAi)* polarity field best matched the experimental results when reducing both the A/P transmission parameter δ_*A*_ and the tail cue α_*T*_ (Fig. S4G). To obtain experimental evidence, we also targeted the planarian homologues of the core PCP-specific component Vang. In contrast to *dvl-1,-2(RNAi)* animals, double knockdown of *Smed-vang-1* and *-2* did not obviously affect tail maintenance (Fig. S6D). However, *vang-1,-2(RNAi)* animals displayed a marked and statistically significant splay increase particularly in central and posterior body regions (Fig. 6G-H), which we could qualitatively reproduce by a reduction of the δ_*A*_ parameter only (Fig. S4F). Altogether, these results therefore indicate that the planarian core PCP pathway is part of the mechanism that orients ciliary rootlets specifically along the A/P axis.

Our model predicts the radial rootlet reorientation upon loss of the A/P component as consequence of the unopposed effect of the Ft/Ds mediated boundary cue (Fig. 6A). Consequently, the concomitant reduction of the margin cue should reduce the magnitude of the ectopic radial component (Fig. 6I). To test this prediction, we targeted *ds* in *dvl-1,-2(RNAi)* animals lacking the A/P polarity component. Knockdown efficiencies in the triple RNAi conditions were similar to the double or single RNAi treatments (Fig. S7A) and triple RNAi animals displayed the tail tip loss of *dvl-1,-2(RNAi)* animals (not shown). Strikingly, the splay pattern of ciliary rootlets was restored to near wild type levels (Fig. 6J-K, Fig. S7B), thus indeed demonstrating the predicted rescue. Together with the fact that we could qualitatively reproduce this effect in our simulations by simply combining the parameter settings of the individual *ds(RNAi)* and *dvl-1,-2(RNAi)* simulations (Fig. S4H), our results establish that the multi-scale coordination of polarity in the planarian ventral epidermis is achieved by the local integration of global A/P and M/L cues (Fig. 7). Furthermore, we identify the planarian core PCP pathway as molecular mediator of the A/P polarity component in our model. Overall, our results therefore establish a mechanistic and quantitative framework for understanding the multi-scale coordination of polarity within the planarian epidermis.

**Figure 7.**
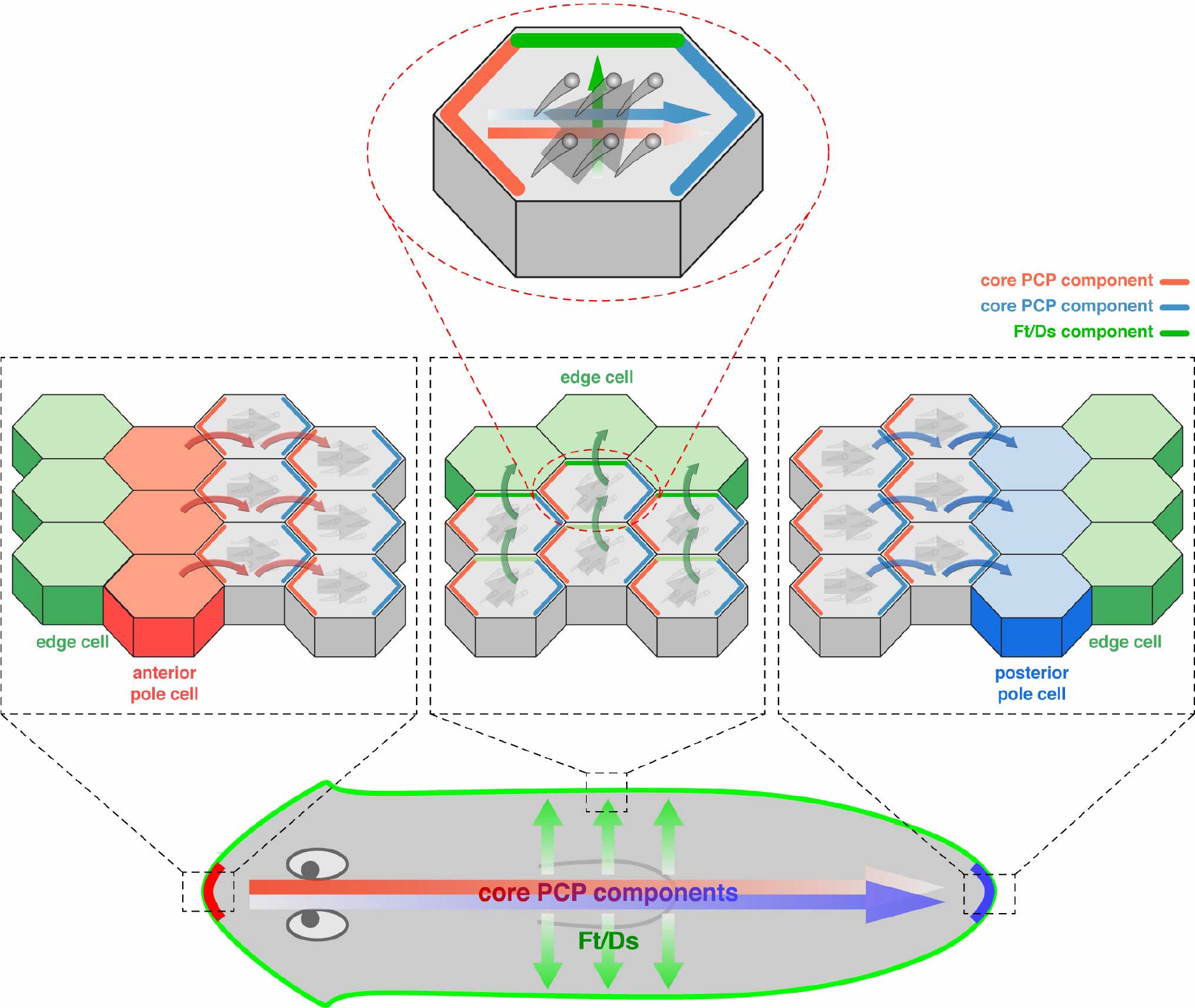
Model of multi-scale coordination of polarity in planarians. See text for details.

## Discussion

Here, we investigate the planar polarization of the planarian ventral epidermis by example of ciliary rootlet orientation. In contrast to the planar polarization of the Drosophila wing disc epithelium that “freezes” after a three day development interval (Adler, 2012), polarization of the planarian epidermis is an inherently dynamic process due to the continuous turnover of its constituent cells (Tu et al., 2015; van Wolfswinkel et al., 2014), similar to the mammalian skin (Devenport et al., 2011). With the ensuing possibility of experimentally repolarizing existing cell fields in the centimeter range, the planarian epidermis epitomizes one of the current key problems in the polarity field: How are consistent polarity patterns established and maintained across scales?

With the extensive mechanistic understanding of core PCP and Ft/Ds-mediated polarization pathways as foundational basis (Adler, 2012; Aw and Devenport, 2017; Butler and Wallingford, 2017; Goodrich and Strutt, 2011; Wallingford, 2010; Yang and Mlodzik, 2015), our results suggest the following model for the multi-scale polarization of the planarian epidermis (Fig. 7). At the cellular level (top panel), the differential subcellular localization of the core PCP (red/blue) and Ft/Ds (green) pathway components establishes independent polarity vectors within epidermal cells. Ciliary rootlets (gray) orient along the sum (gray arrow) of the two vectors. At the organismal level (bottom panel), the anterior and posterior termini of the midline (head and tail poles; red and blue) and the body margin (green) set the core PCP and Ft/Ds vectors of adjacent epidermal cells via the recruitment of specific pathway components to the membrane interfaces. Cell-cell coupling (middle panel) via the established *trans*–interactions of the transmembrane components of the two polarity pathways propagates these boundary effects across the tissue and ultimately to the ciliary rootlets of individual cells. The epidermal polarity field thus emerges as a consequence of spatially restricted boundary effects and their differential superposition (middle panel). Our model generally explains how the experimental induction of ectopic heads or tails can change the polarization of ciliary rootlets. Further, the interplay between the independent A/P and M/L components rationalize the head-tail gradation of the splay in wild type animals (Fig. 2F-H), anterior polarity reversal in “headless” *pbx(RNAi)* animals (Fig. 3H-I) and its rescue upon concomitant knockdown of Ft/Ds components (Fig. 5G-H). Finally, the model explains the restoration of A/P orientation under simultaneous inhibition of Ft/Ds and core PCP (Fig. 6J-K) as restoration of the resultant upon quantitative weakening of both polarity fields (Fig. S4C, G-H). Overall, the model provides a mechanistic framework that links the alignment of subcellular ciliary rootlets to the global landmarks of the planarian body plan.

Our experimental demonstration of separate A/P and M/L polarization components and of core PCP and Ft/Ds as their specific molecular mediators provide a strong biological rationale for the two-field assumption in our mathematical model (Fig. 4), even though the superposition of two fields may seem mathematically non-parsimonious. Indeed, a single polarity field (e.g., direct coupling of H and T to 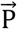 in Fig. 4) is broadly sufficient for transmission of the boundary effects into the epidermis and the qualitative recapitulation of the observed phenotypes (see Supplemental Materials). The strongly margin-enriched expression of *ds* (Fig. 5B) and more uniform core PCP expression (Fig. S6A) would also be compatible with a one-field mechanism, whereby Ft/Ds functions specifically at the body margins to define the boundary conditions for an organismal core PCP-mediated polarity field. Staining protocols and antibodies capable of visualizing the subcellular distribution of Ft/Ds and PCP components will be required to distinguish between the two possibilities. However, our results demonstrate conclusively that ciliary rootlets in the planarian epidermis orient according to polarity inputs from both pathways. To our knowledge, this is the first demonstration of distinct and simultaneous contributions of both major planar polarity pathways to cell polarization outside of *Drosophila*, which therefore establishes the combination of core PCP and Ft/Ds as evolutionarily conserved 2D-polarization module.

The fact that head, tail and body margin orient the A/P and M/L polarization components further demonstrates that these landmarks specify the cellular core PCP and Ft/Ds vectors. This observation is interesting because the same regions are also implicated in the specification of the cardinal body axes of the planarian body plan. Signals expressed by the margin cells participate in midline and D/V patterning (Adell et al., 2009; Gaviño and Reddien, 2011; Gurley et al., 2010), while a small group of cells at the head and tail tips are thought to function as “organizers” of the planarian head and tail (Blassberg et al., 2013; Chen et al., 2013; Oderberg et al., 2017; Vásquez-Doorman and Petersen, 2014; Vogg et al., 2014). These so-called pole cells regenerate in a *pbx*-dependent manner and the headless and tailless animals resulting from *pbx(RNAi)* are one of the lines of evidence that support the critical patterning role of these cells (Blassberg et al., 2013; Chen et al., 2013). Our observation that *pbx* is required for the orienting effect of the head (Fig. 3H-I) implicates the head pole in the orientation of the A/P polarity component. Further, many Wnt signaling components are expressed in form of gene expression gradients along the A/P axis (Gurley et al., 2008; 2010; Lander and Petersen, 2016; Petersen and Reddien, 2009; 2011; Reuter et al., 2015; Scimone et al., 2016; Stückemann et al., 2017) and a tail-to-head gradient of canonical Wnt signaling patterns cell fates along the A/P axis (Stückemann et al., 2017). Therefore, the core PCP-mediated A/P polarization component aligns parallel to the organismal canonical Wnt signaling gradient of planarians, while the orientation of Ft/Ds-mediated M/L polarization component correlates with the margin-enriched expression of *ds* itself. The deployment topology of both pathways is strongly reminiscent of the planar polarization of the *Drosophila* wing disc (Wu et al., 2013). Here, Ft/Ds polarization is generally aligned along the proximodistally graded expression of *ds* and the Golgi-localized kinase *fj* (Ma et al., 2003; Zeidler et al., 2000), and the counter gradients of Ds and Fj expression have been shown to instruct sub-cellular Ft/Ds polarization (Brittle et al., 2012; Clark et al., 1995; Mao et al., 2006; Villano and Katz, 1995). Core PCP in the *Drosophila* wing disk aligns relative to the Wnt-ligand Wingless (Wg) expressing D/V and the Hedgehog-expressing A/P-compartment boundaries (Matakatsu, 2004; Matakatsu and Blair, 2006; 2012; Matis et al., 2014), and Wg production at the D/V compartment boundary (Sagner et al., 2012; Wu et al., 2013) or in individual cell clones (Chu and Sokol, 2016; Wu et al., 2013) dominantly re-polarize neighboring cells. Further, *Drosophila* wing disc compartment boundaries are generally regarded as organizers of the wing disc pattern (Alexandre et al., 2014). Therefore, in both *Drosophila* and planarians, organizer-derived signals or expression gradients “organize” both tissue patterning and polarity vectors and such a dual role of organizing regions is likely to be a general principle of multi-scale polarity coordination.

This view raises the general question of how the organizing regions exert their effect at the mechanistic level. By analogy to the *Drosophila* wing disc, the margin-enriched *ds* expression in planarians might be part of the mechanism responsible for the radial orientation of the M/L component and the identification of the upstream regulators of graded *ds* expression will be important. However, planarian *fj* neither seems to be expressed in a counter gradient to *ds* nor to affect ciliary rootlet polarity upon knockdown (not shown), which may indicate mechanistic differences in polarity propagation between the two species. With respect to A/P polarization, an interesting aspect of our findings is that the independent and opposite polarizing effects of the head and tail (Fig. 3B, E) are both mediated by the core PCP pathway (Fig. 6D). This entails that a single pathway simultaneously specifies two opposing polarity vectors in the tissue. The “domineering non-autonomy” of *fz* versus *vang* mutant clones in the *Drosophila* wing disc provides a conceptually intriguing analogy (Taylor et al., 1998; Vinson and Adler, 1987). Both type of clones dominantly re-orient wing hair polarization vectors of adjacent wild-type cells, but in opposite directions. Mechanistically, this effect reflects the antagonism of Fz and Vang sub-complexes within individual cells, but coupling of Fz/Vang complexes across cell-cell interfaces via the mutual affinity of their extracellular domains (Amonlirdviman, 2005). *fz*^−/−^/wild type (wt) interfaces therefore accumulate Fz, while *vang*^−/−^/wt interfaces accumulate Vang such that propagation of the effect by neighbor-neighbor coupling consequently re-orients adjacent wild type cells into opposite directions. Though speculative in the absence of antibody tools, the recruitment of one of the core PCP sub-complexes by the head and tail in our model (Fig. 7) provides a compelling hypothesis for the independent and opposite polarizing effects of the two landmarks. While the effect of Wnt ligands on intercellular Vang/Fz coupling in *Drosophila* (Wu et al., 2013) and the many Wnt ligands expressed in the planarian tail (Gurley et al., 2010; Petersen and Reddien, 2009) provide a potential mechanism for tail-mediated polarization (see below), molecular activities that target Vang and thus specify the opposite polarization of the PCP vector have so far not been demonstrated in any system. The elucidation of the underlying mechanisms will likely be of general interest, since the alignment of the PCP vector between independent and opposite attractors as in the case of the planarian A/P axis might permit more robust polarization especially of large systems.

Our observations also raise the question of how the landmarks ultimately give rise to tissue- or organism-scale polarity fields. Signaling or gene expression gradients are conceptually appealing in this respect, since the gradient source provides an origin of cell polarization and the slope a long-range orienting principle. Motivated by the experimental evidence mentioned above, many current mathematical models of planar tissue polarization rely on graded differences in component activity or abundance as directional cue (Amonlirdviman, 2005; Burak and Shraiman, 2009; Hazelwood and Hancock, 2013; Yamashita and Michiue, 2016). However, the necessity of signaling gradients remains controversial even in *Drosophila* as *ds* loss of function phenotypes in the wing disc can be largely rescued by uniformly overexpressed *ds* constructs (Matakatsu, 2004; Matakatsu and Blair, 2006; 2012; Matis et al., 2014) and recent results question gradient formation by *Drosophila* Wg (Alexandre et al., 2014). By contrast, the conceptual essence of our model (Fig. 4, 7) is the emergence of the global pattern on basis of local cell-cell interactions, i.e., polarity vector specification at the boundary and subsequent propagation into the cell field by neighbor-neighbor coupling. We implement polarity propagation via a non-directional diffusion term that we have previously shown to emerge from the discrete coupling of each cell’s polarity vector towards the average of its neighbors (Hoffmann et al., 2017). Polarization vectors therefore emerge as a consequence of boundary condition propagation without the need for directional cues. The domineering non-autonomy of *fz* and *vang* mutant clones in the Drosophila wing disc illustrates this principle and it is possible that also the polarizing effects of Wg overexpressing clones (Wu et al., 2013) are due to the propagation of local discontinuities rather than the establishment of a long-range Wnt gradient. The striking long-range effect of topological defects in liquid crystal textures provides a dramatic general illustration of the pattern formation capacity inherent to systems comprised of intrinsically polarized units (Chaikin and Lubensky, 2000) and our model’s predictive capability regarding experimental perturbations further demonstrates the power of such a minimalist view.

However, while the top-down perspective of our continuum model takes the intrinsic polarization of epithelial cells for granted and only considers vector orientation, discrete modeling of the actual PCP machinery captures the self-organized emergence of cell polarization that is inherent to the PCP pathways. The resulting tendency of such systems to get trapped in local energy minima (swirl patterns) is often interpreted as reflecting an inherent need for external polarity coordination (Amonlirdviman, 2005; Burak and Shraiman, 2009; Hazelwood and Hancock, 2013). The fact that both planarian Ft/Ds and core PCP orient parallel to gene expression gradients is noteworthy in this respect. Although we so far failed to detect polarity phenotypes upon knock-down of any of the Fz receptors or Wnt ligands with A/P graded expression, we cannot rule out functional redundancy amongst the 5 Wnt ligands or 3 Fz receptors in this group (Gurley et al., 2008; Stückemann et al., 2017). Further, also our model’s technical implementation of boundary condition propagation via a diffusion term does not a priori rule out a contribution of gradients to the process. Overall, the possibility remains that the many striking gene expression gradients in planarians exert instructive effects on long-range polarization and that the coarse-grain top-down view of our model might consequently unduly simplify the challenge of generating organism-scale patterns. In any case, the dynamic repolarization of existing cell fields at centimeter length scales that we demonstrate here provides a complementary perspective to the emergence of polarity in developmental systems and generally suggests planarians as new model system to study the mechanistic basis of long-range tissue polarization.

## Acknowledgements

We thank Drs. Carl Modes, Frank Jülicher and Suzanne Eaton for comments on the manuscript; Stephanie von Kannen and Heino Andreas for technical support; and Rink lab members past and present for lively discussions. We thank the following MPI-CBG facilities for their support: Antibody Facility, Protein Expression Facility, Light Microscopy Facility, Electron Microscopy Facility and Media Kitchen. This work was supported by the grant “Virtual Planarian” as part of the BMBF eBio initiative and by Max Planck Society funding (J.C.R.).

## Author Contributions

Conceptualization, L.B. and J.C.R.; Methodology, C.B., M.K., S.M., and H.T-K.V.; Investigation, J.A., C.B., M.K., S.M., and H.T-K.V.; Visualization, L.B., M.K., S.M., J.C.R., and H.T-K.V.; Writing – Original Draft, L.B. and J.C.R.; Writing – Review & Editing, L.B., M.K., J.C.R., and H.T-K.V.; Funding Acquisition, J.C.R.; Supervision, J.C.R., L.B., and E.W.G..

## Materials and Methods

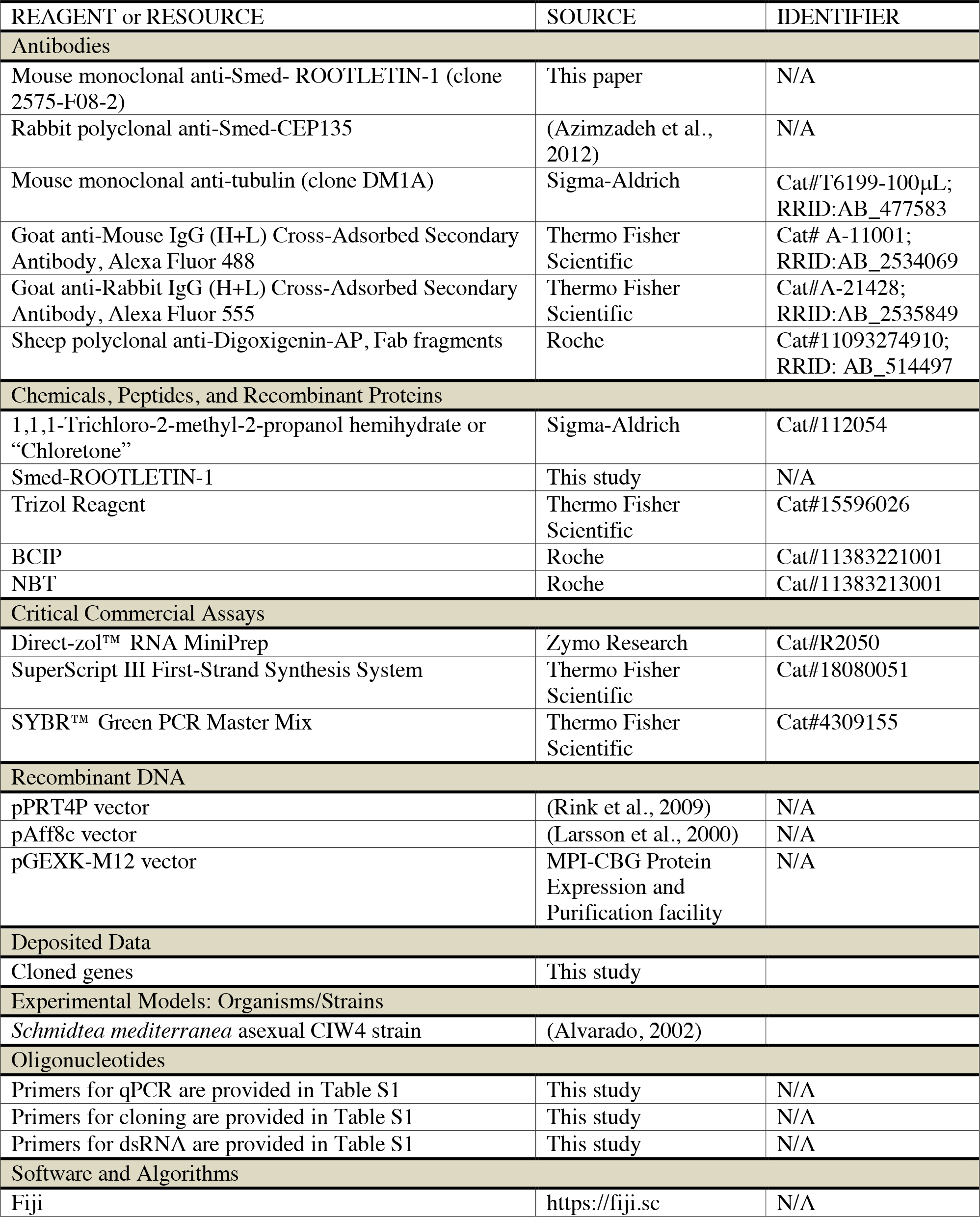
KEY RESOURCES TABLE.

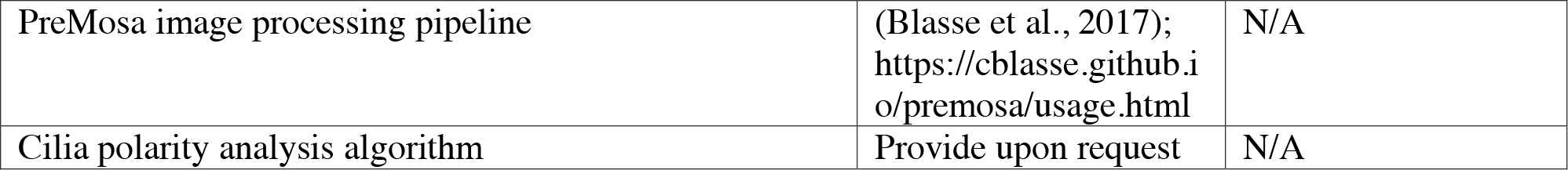

### Planarian Culture

All experiments were performed on asexual *Schmidtea mediterannea* (C4 strain). Animals were maintained at 20°C in 1X Monjuic salts (Cebrià and Newmark, 2005) and fed with calf liver once per week. Animals were starved for at least one week prior to experiments.

### Cloning and RNA-mediated gene silencing

For gene cloning, DNA templates were amplified from cDNA using the primers in Table S1 and inserted into the pPR-T4P vector by ligation-independent cloning (Aslanidis and de Jong, 1990). RNAi of specific target genes was done by feeding liver paste mixed either with dsRNA-expressing *E. coli* (Sánchez Alvarado and Newmark, 1999) or in vitro synthesized dsRNA (Rouhana et al., 2013). RNAi feedings were performed with a two-day gap in between. *β-catenin-1* and *APC(RNAi)* received 3 feedings, all other knock-downs six.

### Recombinant protein and antibody production

The planarian ROOTLETIN-1 antibody was raised against a soluble fragment of the protein. Briefly, the fragment was identified via screening of 64 cDNA fragments resulting from the combinatorial use of 8 regularly spaced forward and reverse primers. Amplified fragments were cloned into the pGEX-M12 vector to add an N-terminal GST-tag, transformed into pRARE-expressing DH5alpha cells, induced with IPTG and the solubility of the specific fragments was assessed by ELISA of culture lysates using standard procedures. For antigen production, the most soluble fragment was retransformed into BL21 competent *E. coli* cells and expressed protein was purified over a Glutathione matrix according to standard procedures (native purification). Monoclonal antibodies were generated at the MPI-CBG Antibody Facility as previously described (Stückemann et al., 2017).

### Electron Microscopy

For ultrastructural analysis by electron microscopy, planarians were transferred into 200 *μ*m-deep flat carriers containing 20% BSA, and high-pressure frozen in an EMPACT2-RTS (Leica, Wetzlar Germany) device. Frozen samples were freeze-substituted in a Leica AFS2 device at −90°C (40 hours), followed by −30°C (8 hours) in a cocktail containing 1% OsO4, 0.1% uranyl acetate, 0.5% glutaraldehyde and 4% ddH2O in acetone and brought to room temperature. The samples were washed in acetone, gradually infiltrated, and flat-embedded in epon-araldite resin. The resin was polymerized at 60°C over the weekend. Small pieces of embedded samples were remounted on a dummy resin block and 100nm-thick sections were cut on a UCT (Leica) ultramicrotome and transferred to copper EM slot grids. The grids were stained for 10 min in 2% uranyl acetate in methanol followed by 2% lead citrate in H2O for 5 min. Images were acquired on a Tecnai 12 (FEI, Eindhoven, The Netherlands) transmission electron microscope equipped with a 2k TVIPS (Gauting, Germany) camera.

### Whole-mount antibody staining

Planarians were anaesthetized in cold 0.2% chloretone in planarian water until completely stretched out, positioned on a filter paper on a cold block with their ventral side up and killed/fixed with ice-cold MeOH for at least 1 hour at −20°C. The samples were then bleached in 6% H2O2 in MeOH overnight under direct light, gradually rehydrated to PBST with 0.1% Triton X-100 (PBSTw0.1%), transferred into reduction solution (1% NP40, 50mM DTT and 0.5% SDS in PBS) for 10 minutes at 37°C, followed by 2×10min washes with PBSTw0.1%. The samples were then blocked for one hour in 10% filtered horse serum in PBSTw0.1%, followed by primary antibody incubation in blocking solution (anti-SMED-ROOTLETIN-1 1:500, and anti-SMED-CEP135 1:500) overnight at 4°C. The samples were washed 6-8 times for 4 hours in PBSTw0.1%, and then incubated in secondary antibody in blocking solution (both secondary antibodies were used at 1:500) for 4 hours at room temperature (or overnight at 4°C). Stained samples were washed 4-6 times for 2 hours in PBSTw0.1%, and then mounted in 80% glycerol (prepared in 10mM Tris pH 8.5).

### In situ hybridization

Whole-mount in situ hybridization was performed as previously described (King and Newmark, 2013; Pearson et al., 2009). Following NBT/BCIP development, animals were mounted in mounting media containing 75% glycerol and 2M urea.

### qPCR

Total RNA from RNAi animals was extracted by mechanical homogenization in Trizol (Invitrogen) and purified using Direct-zol RNA MiniPrep kit (Zymo Research). cDNA was synthesized using SuperScript III reverse transcriptase (Invitrogen). qPCR was performed using SYBR Green PCR Master Mix (Applied Biosystems). qPCR primer sequences are provided in Table S1. Relative RNA abundance was calculated using the delta-Ct method after verification of primer amplification efficiency.

### Light microscopy and image analysis

All imaging was performed on a spinning disk Zeiss microscope equipped with a 40x Oil objective. Images were acquired as a tiled series of individual Z-stacks covering the entire ventral epidermis of the specimens. The resulting 3D image mosaic was processed using the image processing pipeline PreMosa (Blasse et al., 2017), which extracts the signal from the epidermal cells (on basis of the ROOTLETIN-1 and CEP135 channels) and tiles individual images. The PreMosa output is a 2D representation of the ventral epidermis, which was used as basis for all subsequent analyses.

For rootlet detection, the rootlet signal was first reduced to present its major axis by applying a Difference-of-Gaussian filter (σ_1_ = 0.5, σ_2_ = 5) (Sonka et al., 2007). To segment the rootlets in the enhanced images, an adaptive thresholding approach was applied using a local threshold being determined by the Ostu method (Otsu, 1979). To remove false positives or unresolved clusters of ciliary rootlets, connected components that do not fit the expected size of a ciliary rootlet (8-10 pixels under our imaging conditions) were filtered out. The same approach was applied for segmenting the basal bodies.

To obtain the rootlet orientation, an ellipse was fitted to each detected component. The orientation and length of the major and minor axes are given by the eigenvectors and eigenvalues of the resulting covariance matrix. All detected components with major axes (eigenvector of the largest eigenvalue) < 2 than the minor axis (eigenvector of the second largest eigenvalue) were filtered out, since orientation can only be meaningfully determined for elongated ROOTLETIN-1 positive structures. For all remaining rootlets, the largest eigenvector presents the major axis and, thus, orientation. To determine the sign of the orientation within the interval [0°,360°], the asymmetric position of the CEP135-positive basal body at the base of the rootlet was used. Unique assignments of the closest basal body to each ciliary rootlet were made using a formulation of a bipartite matching problem (See Supplementary materials for details).

To approximate cell outlines, we used Voronoi tessellation with the regularly spaced epidermal nuclei as starting points for partitioning the image into polygons (Szeliski, 2011). Nuclei were detected using the Fiji plugin *Descriptor-based Registration (2D/3D)* (Schindelin et al., 2012; Smith et al., 2015).

### Mathematical methods for ciliary rootlet statistics

Data of individual ciliary rootlet angles were treated as normal vectors and averaged within boxes of a 150×150 pixel grid, approximately representing the area of one to two cells under our imaging conditions. The resulting average vector per box was normalized to unit length. Data from cilia close to the tissue border and the pharynx were excluded. The resulting vector maps were used for all statistical analyses. For visualization of vector maps in the figures, coarser grids were used in order to obtain a visible vector size across different animal sizes (grid size was adjusted to specimen size). Graphs quantifying the splay of rootlet vectors against the A/P axis were calculated by averaging vectors along stripes in horizontal or vertical directions using a coordinate system that was aligned to the long and short main axes of each individual animal, i.e. the best-fitting ellipsoid to its shape, and scaling position by axis length resulting in relative position values from 0 to 1.

## Supplemental material for mathematical modeling

### Details on the model equations

Two vector fields 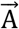 and 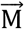 are defined on a two-dimensional domain approximating the planarian body shape, rendering the precise value of 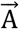 and 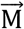 a function of the spatial position 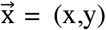. These two fields represent two polarity components contributing independently to the observable ciliary rootlet polarity and are generated by organizing regions at the head, tail and body margin.

For the geometric definition of the head, the tail and the body margin, we use the three static functions H(x,y), T(x,y) and 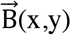 that are markers for the head (red), the tail (blue) and the entire boundary (green curve in Fig. 4A), respectively. H is set 1 where a head is located (at the anterior side in the wildtype) and 0 everywhere else. Analogously, T is set 1 where the tail is located (at the posterior side in the wild type) and 0 everywhere else. B is a vector of length 0 in the interior of the animal and of length 1 on the boundary pointing outwards and perpendicular to the boundary of the computation domain.

The spatiotemporal dynamics of 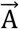 and 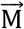 are given by two partial differential equations (PDE) in Fig. 4B. The operator B̃ is defined here as 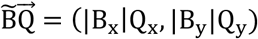 with 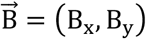 and 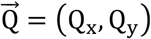. The terms in these equations can be understood as follows. The first term (grey background) represents the action of the organizing regions at the head, the tail and the boundary. It generates nonzero values for 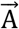 and 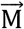 at certain locations at the boundary. For 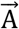 this happens at the head (encoded by H) and the tail (encoded by T). For 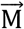 this happens at the whole boundary (encoded by 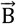). The areas of H and T are chosen 5 lattice units wide, corresponding to 1% of the body length, and as long as 10% of the body width, as indicated by the colored areas in all simulation result figures. The term ~α is a source term and the term ~β is a degradation term, both terms are only active on the respective boundary area. If no other terms in the PDE were present and the dynamics reached the steady state, 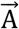 would have magnitude α_*H*_/*β*_*H*_ at H and point inwards 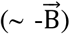 and 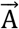 would have magnitude α_*T*_/*β*_*T*_ at T and point outwards 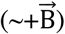. Similarly, if no other terms were present, 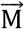 would have magnitude α_*B*_/*β*_*B*_ at the boundary 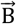 and point outwards. Therefore, we can adjust the strength of the organizing regions at the boundary, the head and the tail by adjusting the α and the β parameters. In the presence of the other terms the values of 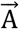 and 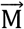 at the boundaries will change slightly since the described dynamics at the boundary is not equivalent to classic Dirichlet boundary conditions but allows us to model the dynamic adaptation of the boundary states in response to perturbed cues like the reduction of strength (~α). Moreover, our dynamic and more flexible boundary effect can balance between boundary preference and bulk dynamics with few uniform parameters.

The second term (green background) represents degradation of the vector magnitude due to stochasticity and cell turnover, with subsystem-specific rates γ_A_ and γ_M_. The third term (blue background) is a spatially averaging Laplacian with strength D_A_ and D_M_ representing the neighbor-neighbor coupling as used previously in continuum models of fly wing polarity (Merkel et al., 2014). The Laplacian acts component-wise on the vectors. Numerically, a square lattice and no flux boundary conditions around the body margin are used.

To obtain the polarity field 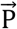, the two fields 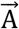 and 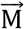 are superimposed on each other by a simple linear combination and then normalized to modulus 1 for non-vanishing superposition. The parameters δ_A_ and δ_M_ represent the responsiveness of the cells to the 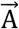 and 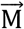 fields. These parameters range from 0 to 1 where 1 represents complete responsiveness and 0 complete unresponsiveness.

### Simulation details

The equations were simulated using Morpheus (Starruy et al., 2014). The simulations were performed on a domain having an approximate planarian body shape on a 160×50 grid with Δx = 0.5. A forward Euler method with Δt = 0.05 was used for integrating the equations. Except for the *pbx(RNAi)* and the *pbx/ds(RNAi)* simulations, the equations were integrated until steady-state was reached.

### Parameter estimations

#### Control condition

For the wild type in the model, we set δ_A_ and δ_M_ to 1, representing complete responsiveness to the polarity components 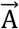 and 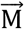 and fit the other parameter values as follows. The approximately linear decrease of splay along the anterior-posterior axis indicates that the degradation terms are small and that D_A_ is large but D_M_ is relatively smaller. Because the strengths of the head, tail and boundary organizing regions are set by the respective ratios α/β, we fix all β parameters to 1.0 and search for values of the α parameters that match the experimentally measured splay profile. Again, this task is made easier by the gradual change in splay along the anterior-posterior axis. By symmetry, in the center of the animal the x projection of 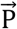 equals 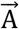 and the y projection equals 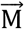. Hence, in the center of the animal the tangent of the splay equals 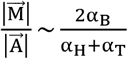. Analogously, similar relations can be found at other positions of the animal. Inserting experimentally measured splay angles thereby informed our choice of the α parameters. Estimates for the a parameters were refined by manual iteration until the observed splay profile was
matched closely (Fig. S4A). As it turns out small changes in these parameters do not affect the vector field 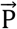 significantly. All parameter choices are shown in Table S2.

#### β-catenin-1(RNAi) *and* APC(RNAi)

The parameter values for simulating the *β-catenin-1(RNAi)* and *APC(RNAi)* cases are exactly the same as for the Control. The only differences for the simulations are the T and H functions. For *β-catenin-1(RNAi)* we set T^bcat^ = 0 everywhere (there is no tail present) and H^bcat^ = H^ctrl^ + T^ctrl^ (there is another head where the tail was before). Similarly, we set H^apc^ = 0 everywhere (there is no head present) and T^apc^ = H^ctrl^ + T^ctrl^ (there is another tail where the head was before). Fig. 4D, E show simulation results that match the observed polarity fields.

#### Remaining RNAi experiments

The molecular details of the *pbx(RNAi)* experiments suggests inactivated head and tail organizers, that means we should have 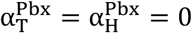. This however, would mean that the A/P polarity component completely disappears, which is not what we observe. Therefore, we simulate the *pbx(RNAi)* condition by starting with the Control steady-state, switch off head and tail by setting 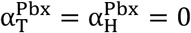 and assume that the observed polarity pattern upon *pbx(RNAi)* corresponds to a transient state at a certain time t when the A/P field 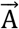 has not yet completely decayed. In the simulations we used t = 2.5. In this case a very good fit to the observations is achieved (Fig. 4F and Fig. S4B).

All following simulations are carried out until the steady state is reached. For most of the remaining RNAi experiments all parameters are taken unchanged from the Control case and only the δ_A_ and δ_M_ parameters are adjusted. In the *ds(RNAi)* and *ft-1,-2(RNAi)* conditions, smaller splay is observed than in the Control, therefore we fix δ_A_ = 1 for both cases and manually adjust δ_M_ until the splay in the simulations is close to the experiments. As a result we find 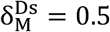 and 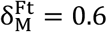. This model fit with a reduced role of the 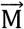 is in agreement with the putative role of Ds and Ft in propagating the M/L cue from the body margin. For the results see Fig. S4C-D.

The *pbx/ds* double RNAi condition is simulated by simply combining the *pbx(RNAi)* parameters with the *ds(RNAi)* parameters and not fitting any parameter. This means we again determine the fields 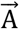 and 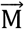 at a time t = 2.5 after inactivating head and tail and use 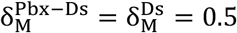. With these parameters a very good agreement is found between experiments and simulations (Fig. S4E).

We proceed similarly for the *vang-1,-2(RNAi)* conditions. As in these animals larger splay is observed than in the Control we fix δ_M_ = 1 and find by manual fitting 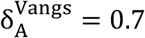, in agreement with a role for Vangs in propagating the A/P cue from the head and tail (Fig. S4F).

For the Dvl case this procedure of reducing 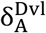 does not result in satisfying fits. The splay is either too large in the anterior region or too small in the posterior region. However, the situation changes after we switch off the tail organizer by setting 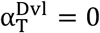, which is in agreement with a dual role for Dvls in canonical PCP propagating the A/P cue and in tail specification. With the inactivated tail, 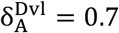 yields a good fit to the observed splay profile (Fig. S4G), keeping the 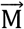 field unperturbed (δ_M_ = 1). Note, if we reduce the cell-cell coupling strength D_A_ as motivated by a possible role of Dvl in Frz-Stbm transcellular complex formation, then splay approaches 90 degrees away from the anterior and posterior poles. While the reduction of D_A_ alone does not reproduce the experimentally observed splay profile (about 65 degrees in the trunk region) as well as the above parameter set, a moderate D_A_ reduction could complement the above parameter set and yield a similar model result (data not shown). Importantly, combining the settings for *dvl-1,-2(RNAi)* and *ds(RNAi)* provides the parameters for the *dvl-1,-2/ds* triple RNAi condition, i.e. 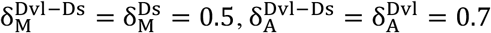 and 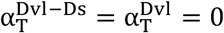. This fit-free model prediction of the triple RNAi condition matches well with the observed splay profile (Fig. S4H).

### Discussion of alternatives and extensions of our boundary model

Our integrated mathematical modeling results support the collective sufficiency of local boundary conditions plus neighbor coupling for explaining the multi-scale polarization of the planarian epidermis. Specifically, by means of iterative model extension starting from published core models (Amonlirdviman, 2005; Burak and Shraiman, 2009; Merkel et al., 2014) and comparison of model performances among alternative mathematical models, we identified one model as shown in Fig. 4 that reproduces the experimental data quantitatively (Fig. S4) and only uses one new concept: the superposition of two separate vectorial polarity cues, termed 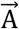 and 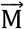 for A/P cue and M/L cue, respectively. Correspondingly, this superposition model only possesses few degrees of freedom which have been determined by fitting a subset of the experimental observations (see above). We chose to adjust the responsiveness parameters δ_A_ and δ_M_ of 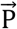 to 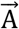 and 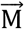 but similar results can be obtained if the amplitudes of the 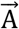 and 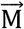 fields are adjusted through α since field amplitude and response are cumulative factors. The calibrated model then correctly predicted two independent experimental conditions of multiple simultaneous RNAi perturbations (Fig. S4E, H).

Note, an interesting and simpler alternative model does not reproduce the observed spatial dependencies of the splay effect as well as our model: a single vector field 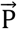 under the influence of head, tail and lateral boundary conditions would be able to qualitatively reproduce some of the experimental observations, but differs quantitatively for some of the experimental conditions including the Control. Another model variant could specify fixed Dirichlet boundary conditions along the body margin, thereby replacing the α and β terms in Fig. 4B, but would generate discontinuities at the edges of the organizers and constant splay along the trunk, which are both contradicting our observations. There are additional known processes which the presented model does not explicitly take into account, including epithelial cell turnover, heterogeneity of cell types (incl. gland cells) and neighbor-neighbor coupling strength or the presence of additional boundary conditions around the pharynx. To test the robustness of our model results against such additional components, we considered a stochastic model extension by adding δ-correlated random variables of low magnitude to the right hand sides of the equations for 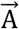 and 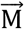 (Fig. 4B) and found a local and random modulation of all polarity patterns that mimics the noise in the observed data but otherwise lets the model results unchanged (data not shown). Epithelial cell turnover can implicitly be accounted for by the choice of the neighbor-neighbor coupling strength, where higher turnover rate reduces the effective coupling strength, and need not be considered explicitly as theoretically shown (Hoffmann et al., 2017).

In the presented model, head and tail instruct polarity through boundary conditions which is found to be sufficient and minimal. An extended model could replace the boundary conditions for 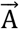 and 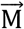 fields by three ligand concentration fields, forming gradients from sources at head, tail and lateral boundary, diffusion and decay within the field, and local orientation of the vector fields towards the direction of the ligand gradients (Burak and Shraiman, 2009; Rappel and Edelstein-Keshet, 2017). If the characteristic decay lengths of the resulting gradients, proportional to the square root of the ratio of ligand diffusion and decay, are small compared to the animal body size then such extended models will, at the scale of the animal, show identical results as presented here.

Hence, on the one hand, multiple molecular mechanisms at or near the boundaries can effectively be described by our model. On the other hand, the simplicity of our model with just polarity decay and neighbor coupling inside the spatial domain does not account for spontaneous pattern formation like swirls from random initial conditions (Burak and Shraiman, 2009). Our model can be extended by adding a normalizing flux 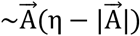 or 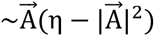 to the right hand side of the model equation for 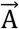 (Fig. 4B) and swirl patterns become possible starting from random initial conditions in the absence of boundary effects. The choice of η then adjusts the modulus of 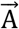 and contributes to the balance of the polarity components 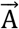 and 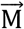. Further, our model does not incorporate any mechanical influence on polarity such as resulting from shear flow of cells relative to each other as observed in the fly wing (Aigouy et al., 2010; Eaton and Jülicher, 2011; Merkel et al., 2014). While shear flows result from tissue growth and hinge contraction next to the fly wing primordium, there are no such forces acting on the planarian epithelium. However, cell extrusion and cell incorporation at different locations, as long as such random processes do not lead to uniform average rates, would generate shear flows within the epithelium and would turn individual cells relative to the resting reference frame of the tissue boundary, thereby affecting polarity patterns. Further experiments quantifying the spatial dependency of cell extrusion and cell incorporation rates will be necessary to evaluate the magnitude of shear flows. Our model could then easily be extended for both fields 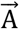 and 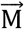 by additional flow terms as in the models of wing hair polarity (Aigouy et al., 2010; Merkel et al., 2014). Altogether, our model is simple, reproduces the studied planarian polarity patterns, has correctly predicted two additional combination conditions and is extensible to include the effects of fly wing patterning.

## Extended explanation of rootlet orientation quantification

Since polarities comprise a directed orientation within the interval [0°,360°], it is still necessary to still determine the correct sign of the estimated orientation. From the literature, it is known, that the basal body appears at the base of the rootlets, thus the position of the basal bodies allows the determination of the unknown directionality. We utilize a formulation of a bipartite matching problem to uniquely assign the closest basal body to each ciliary rootlet. Let R be the set of ciliary rootlets and B the set of basal bodies, then a matching m = (V,E) can be defined as a weighted bipartite graph with:

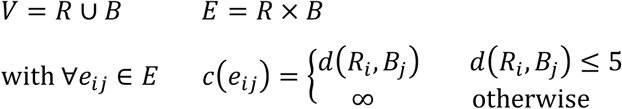

For each edge *e*_ij_ ∈ *E*, the cost *c*(*e*_ij_) is defined as the Euclidean distance between the center of the basal body *B*_j_ and the closest tip of the rootlet *R*_i_. To reduce the size of the matching problem and to prevent implausible assignments, only edges with a distance less or equal than 5 are considered. Using a greedy algorithm to compute a matching reveals a solution that locally minimizes the total costs of used edges and rather leaves a rootlet or basal body unmatched than assigning them to the wrong matching partner. Even if this does not obtain a global optimal solution, the majority of basal bodies are correctly matched. For each ciliary rootlet, the solution reveals either an exclusion from the analysis, if no reasonable assignment could be determined, or a unique basal body that localizes near one of its tips. The position of a matched basal body finally enables the determination of a direction of the previously estimated rootlet orientation. With that, it is possible to assign a polarity p = P(θ) ∈ {1, …, 8} to the ciliary rootlet being elongated and having a basal body near its tip.

To infer the polarity for entire cells or tissues, a cell segmentation is required, which yields the area of each cell within the epithelium. Since it was not possible to visualize cell membranes uniformly in our immunostaining protocol, the cell segmentation is determined from a Voronoi tessellation having the cell nuclei as starting points. To detect all cell nuclei in the 2D image, we apply the Fiji plugin *Descriptor-based Registration (2D/3D)* (Schindelin et al., 2012; Smith et al., 2015). This plugin is a tool to register two images by matching two point clouds whereby the used points are identified by a Difference-of-Gaussian detector accepting only blob-like structures of a predefined size. Although it is designed to register two images, we exploit it to detect cell nuclei, which also yield roundish structures of a specific size. The detected positions serve as starting points to partition the image into polygons using a Voronoi diagram (Szeliski, 2011). Since we could observe, that the cell nuclei locate exactly in the center of the cell, the Voronoi tessellation is a reasonable approach to estimate the cell boundaries.

To summarize the rootlet polarities of each cell being expressed by both an angle θ ∈ [0°,360°] and a polarity p = P(θ) ∈ {1, …, 8}, we determine the cell polarity p_C_ via majority voting:

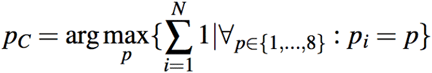

The analysis revealed that each cell features on average 30 rootlets. Thus, if a cell deviates significantly from that, then there must be a mistake in the cell segmentation clustering or splitting correct cells. To exclude these cells from the analysis, we reject cells featuring less or more than the allowed rootlet number. Currently, we accept cells having 10-70 rootlets. All remaining cells feature a reasonable rootlet number, which is summarized to a common cell polarity.

## Supplemental Figures

**Fig. S1:**
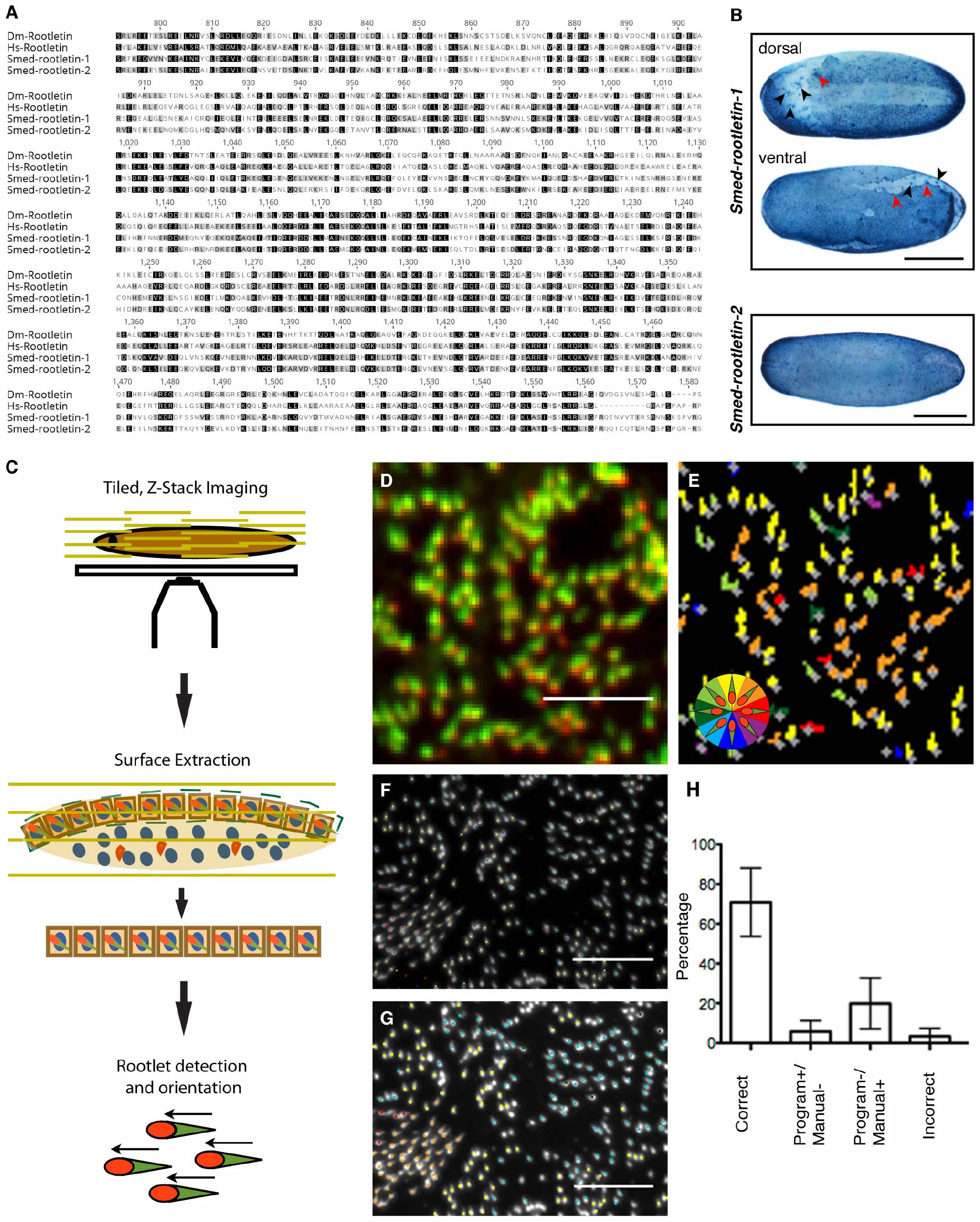
Ciliary rootlet visualization and analysis. **A.** Amino acid sequence alignment (Clustal-W) of an 800 amino acid fragment of *D. melanogaster* (Dm) Rootletin, *H. sapiens* (Hs) Rootletin, *S. mediterranea* (Smed) ROOTLETIN-1 and ROOTLETIN-2. Black: Conserved, gray: similar residues (Blosum62 as score matrix). **B.** Whole mount gene expression patterns of *Smed-Rootletin-1* and *−2* by *in situ* hybridization. Red arrowhead: Tear in the epidermis that highlights epidermal expression. Black arrowhead: Protonephridial expression. Scale bar: 500 μm. **C.** Cartoon illustrating the multi-scale ciliary rootlet analysis pipeline. First, spinning disk confocal imaging of entire whole mount specimens by tiled Z-stacks achieves global visualization of micron-length ciliary structures. Second, computational extraction of the curved epidermal layer from the stitched Z-stacks using a customized surface extraction algorithm ((Blasse et al., 2017); see Materials and Methods for further detail information). Third, automated rootlet extraction and analysis using custom-developed algorithms (see Materials and Methods for further information). **D.** High-magnification view of rootlets (ROOTLETIN-1 (green) and CEP135 (red) antibody staining). **E.** Same cell field as in (D) with grey crosses designating detected rootlets and colored extensions rootlet orientation as per the indicated color wheel. Comparisons between D and E demonstrate the generally high sensitivity and specificity of the algorithm with respect to rootlet detection and orientation. **F-H.** Quantitative validation of algorithm performance. Automatic (F) and manual ground truth annotation (H) of a representative cell field in the ROOTLETIN-1 channel. Dots designate ciliary rootlets co-labelled by CEP135 staining (not shown) and color encodes orientation as per the color wheel in E. H: Bar graph comparison of automatic versus manual annotations. “correct” designates annotations within +/− 45°, “incorrect” > +/− 45°,and the remaining two categories events scored by one of the two methods only. Overall, the comparison demonstrates correct directionality assignments for ~ 70% of detected rootlets, with false negatives (i.e., undetected rootlets) according for the majority of errors. Wrong directionality assignments accounted for < 5% of cases.

**Fig. S2:**
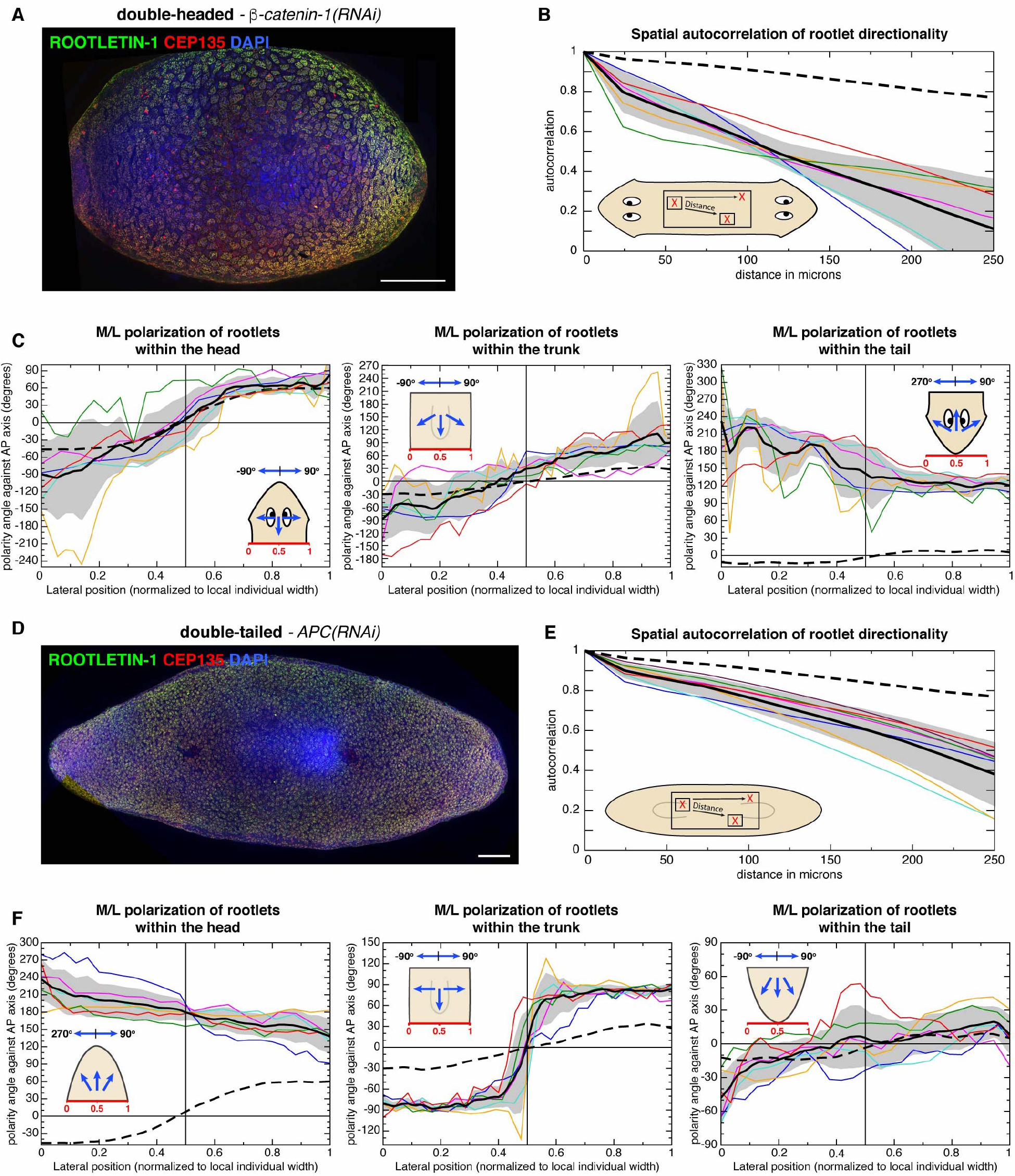
Head and tail as independent organizers of tissue polarity. **A.** Stitched and surface extracted high-resolution image of the entire ventral epidermis of a representative double-headed *β-catenin-1(RNAi)* specimen 14 days post amputation. Anti-ROOTLETIN-1 (green), anti-CEP135 (red) and nuclear staining (Hoechst, blue). Scale bar: 500 μm. **B.** Spatial autocorrelation analysis of rootlet directionality in 6 double-headed specimens, comparing directional coherence within square of 150×150 pixels. **C.** M/L polarization component in 6 double-headed *β-catenin-1(RNAi)* specimens. Colored lines trace the averaged rootlet orientation parallel to the midline in the head (left), trunk (middle) and “tail” (right). Traces were normalized by width. **D.** Stitched and surface extracted high-resolution image of the entire ventral epidermis of a representative double-tailed *APC(RNAi)* specimen 14 days post amputation. Anti-ROOTLETIN-1 (green), anti-CEP135 (red) and nuclear staining (Hoechst, blue). Scale bar: 500 μm. **E.** Spatial autocorrelation analysis of rootlet directionality in 6 double-tailed specimens, comparing directional coherence within square of 150×150 pixels. **F.** M/L polarization component in 6 double-tailed *APC(RNAi)* specimens. Colored lines trace the averaged rootlet orientation parallel to the midline in the head (left), trunk (middle) and “tail” (right). Traces were normalized by width. B, C, E, F: Colored lines represent individual specimens. The solid black line and the grey shading indicate mean plus/minus standard deviation of 6 animals. 0° designates head to tail orientation, +/− 90° orthogonal orientations and cartoons illustrate measured rootlet orientations.

**Fig. S3:**
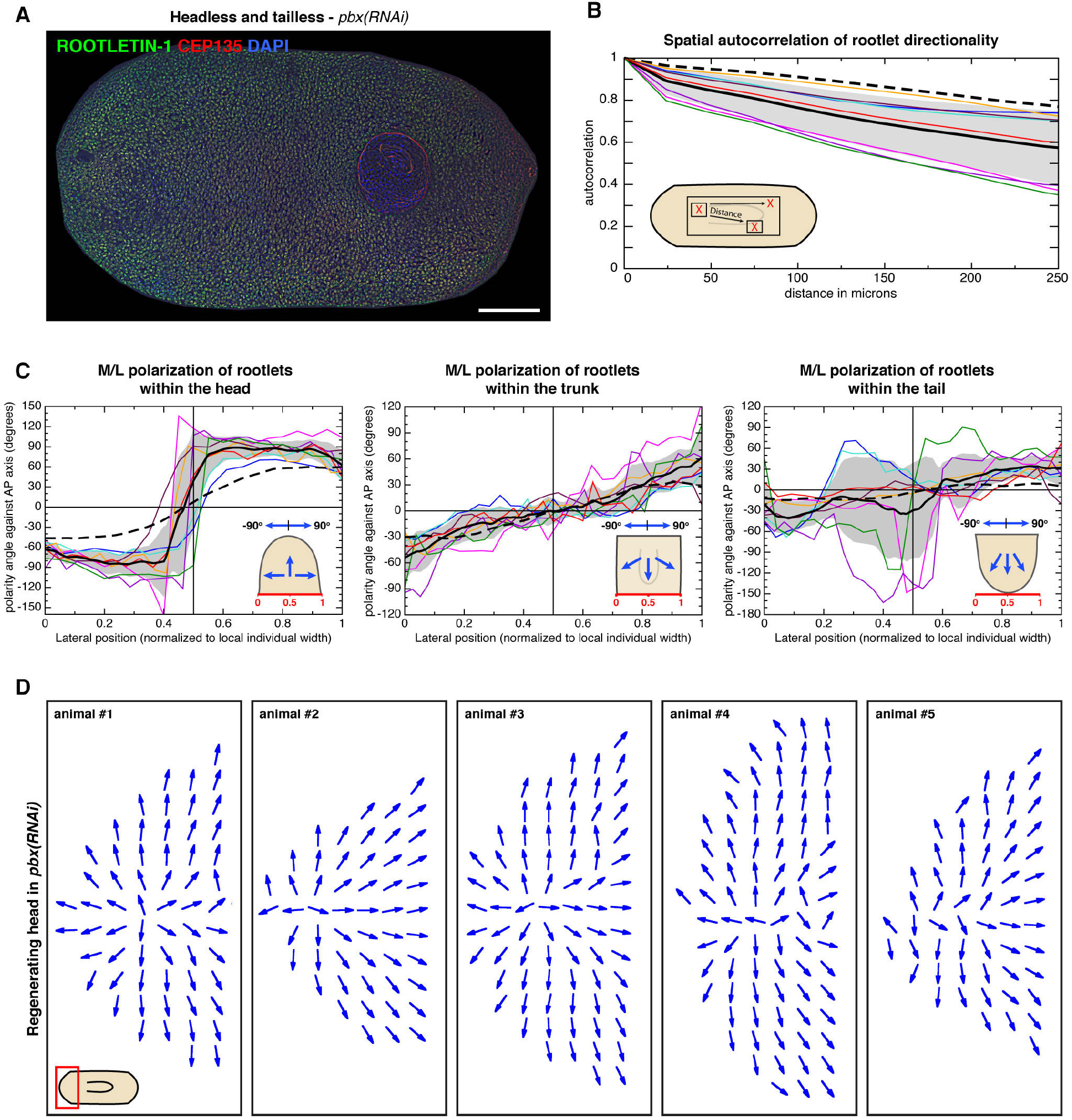
The body margin as organizer of tissue polarity. **A.** Stitched and surface extracted high-resolution image of the entire ventral epidermis of a representative head/tailless *pbx(RNAi)* specimen 14 days post amputation. Anti-ROOTLETIN-1 (green), anti-CEP135 (red) and nuclear staining (Hoechst, blue). Scale bar: 500 μm. **B.** Spatial autocorrelation analysis of rootlet directionality in 6 head/tailless specimens, comparing directional coherence within square of 150×150 pixels. **C.** M/L polarization component in 6 headless and tailless *pbx(RNAi)* specimens. Colored lines trace the averaged rootlet orientation parallel to the midline in the head (left), trunk (middle) and “tail” (right). Traces were normalized by width. **D.** Vector field representation of the average ciliary rootlet orientation within 100 μm bins in the anterior region of 5 representative head/tailless *pbx(RNAi)* specimens 14 days post amputation. B, C: Colored lines represent individual specimens. The solid black line and grey shading indicate mean plus/minus standard deviation, respectively. The dashed line represents the average of 6 wild type specimens. 0° designates head to tail orientation, +/− 90° orthogonal orientations and cartoons illustrate measured rootlet orientations.

**Fig. S4:**
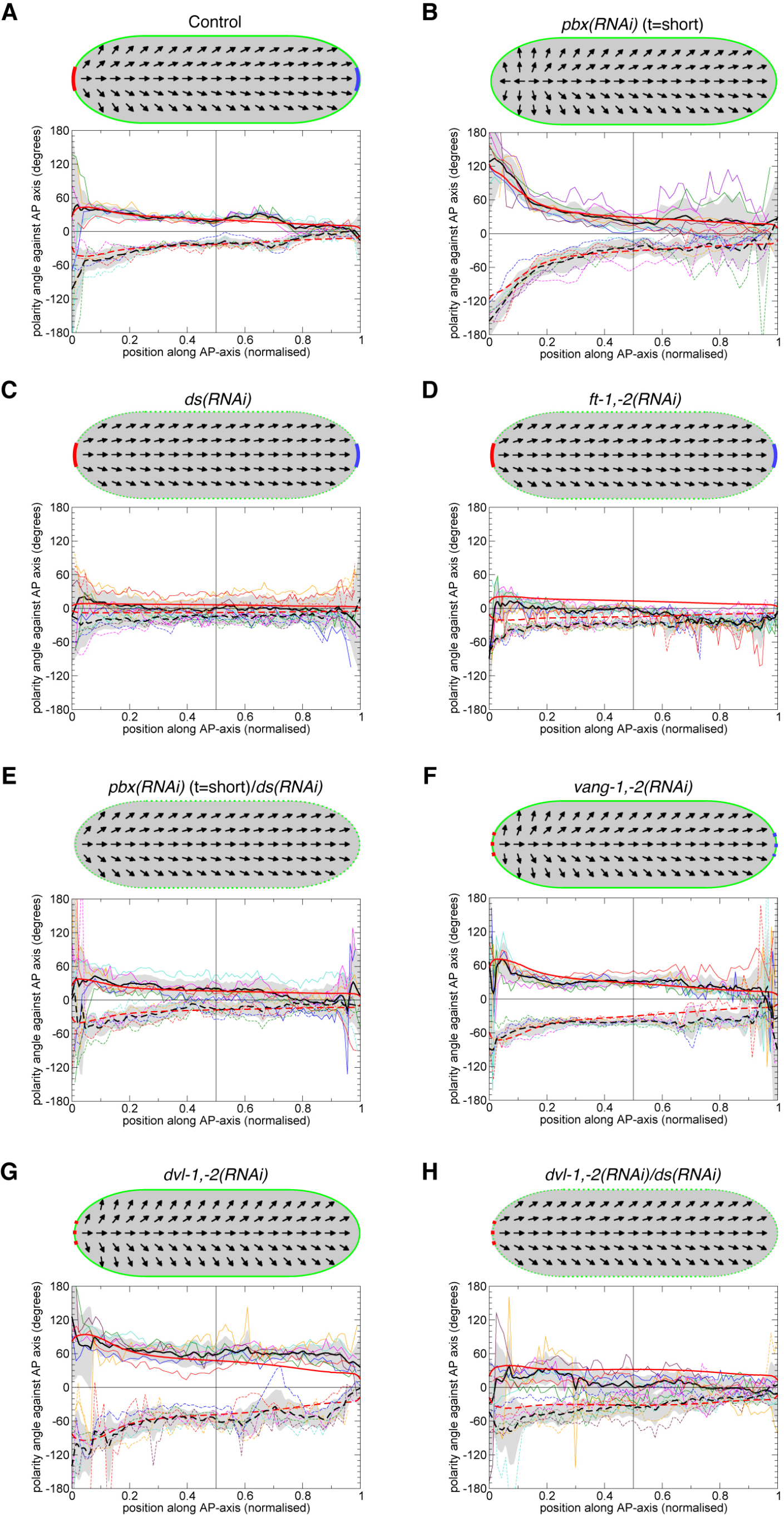
Simulations and quantitative comparison to experimental data. **A-G.** Simulated 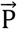 fields as vector plots of the indicated experimental conditions. **A’-G’.** Average splay angle as function of A/P position for the simulation (thick red) overlaid with experimentally measured traces of the respective experimental condition (thin colored lines) and their average (thick black line). The gray band denotes plus/minus standard deviation. **A’.** Parameter values for model simulations (Table S2) were obtained by minimizing the distance between model (thick red) and data (thick black) curves. Note, the model is inherently left (dashed) versus right (solid) symmetric, while the experimental data may indicate slight discrepancies. **B’, C’, E’, F’.** Experimental fitted by tuning only a single parameter value as motivated by each condition. **D’.** Simulation of the *pbx/ds* double RNAi condition as combination of the parameter settings of the two single RNAi conditions. **G’.** Qualitative agreement between simulated (red) and average observed (black) ciliary rootlet orientation for *dvl-1/dvl-2/ds* triple RNAi by a combination of the two individual parameter settings.

**Fig. S5:**
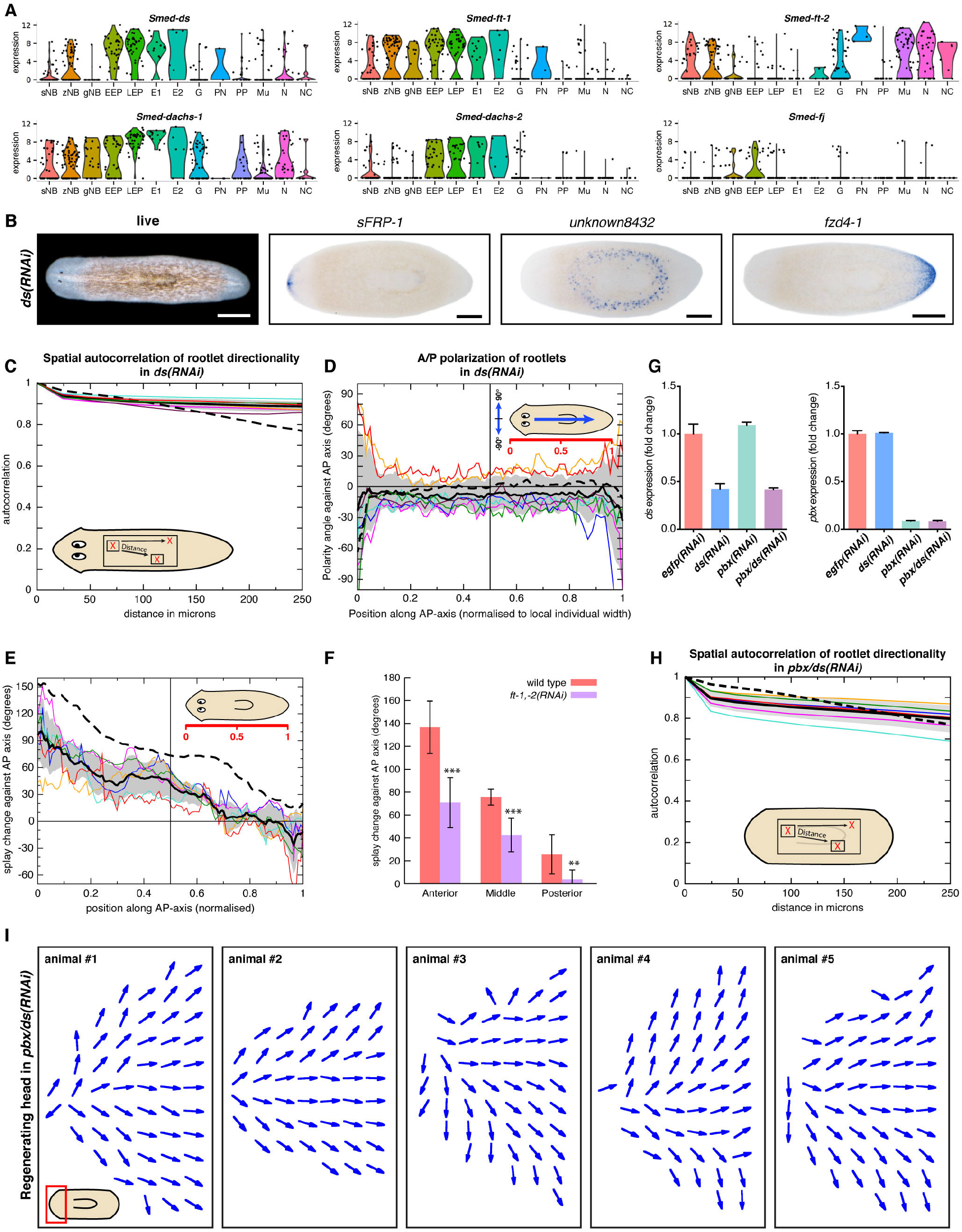
Ft/Ds is required for the ML polarity component. **A.** Single cell gene expression profiles of Ft/Ds pathway components in different cell types, generated using the Planarian SCS online resource (Wurtzel et al., 2015). sNB: sigma neoblast; zNB: zeta neoblast; gNB: gamma neoblast; EEP: early epidermal progenitors; LEP: late epidermal progenitors; E1: epidermis I; E2: epidermis 2; Gut: gut; PN: protonephridia; PP: parapharyngeal; Mu: muscle; N: neural; N-C: neural (ciliated). **B.** Body plan polarity landmarks in representative *ds(RNAi)* specimens 14 days post amputation. Live image (left), whole mount in situ hybridization gene expression patterns of head, trunk and tail markers to the right. Scale bar: 500 μm. Normal marker expression indicates absence of overt regeneration polarity phenotypes. **C.** Spatial autocorrelation analysis of rootlet directionality in 7 intact *ds(RNAi)* specimens, comparing directional coherence within square of 150×150 pixels. **D.** A/P-polarization component in intact *ds(RNAi)* specimens: Lines trace the averaged rootlet orientation orthogonal to the midline in 7 intact *ds(RNAi)* specimens, traces were normalized by length. Colored lines represent individual specimens. The solid black line and grey shading indicate mean plus/minus standard deviation of 7 animals, respectively. 0° designates head to tail orientation, +/ - 90° orthogonal orientations and cartoons illustrate measured rootlet orientations. **E.** Splay angle change along the A/P axis under *ft-1,-2(RNAi)*. Colored lines represent 6 individual specimens. The solid black line and grey shading indicate mean plus/minus standard deviation, respectively. The dashed line represents the mean of 6 wild type specimens. 0° designates uniform A/P orientation, 180° M/L orientation. **F.** Quantitative comparison of splay angle changes between *ft-1,-2(RNAi)* and wild type, based on E. Splay angles of 6 specimens each were averaged within the head, trunk and tail regions. * p-value < 0.05; ** p-value < 0.01; *** p-value < 0.001. **G.** Gene expression levels of *pbx* (left) and *ds* (right) by qPCR under the indicated single- and double(RNAi) conditions. **H.** Spatial autocorrelation analysis of rootlet directionality in 6 *pbx/ds(RNAi)* specimens 14 days post amputation, comparing directional coherence within square of 150×150 pixels. **I.** Vector field representation of the average ciliary rootlet orientation within 100 μm bins in the anterior region of 5 representative head/tailless *pbx/ds(RNAi)* specimens 14 days post amputation.

**Fig. S6:**
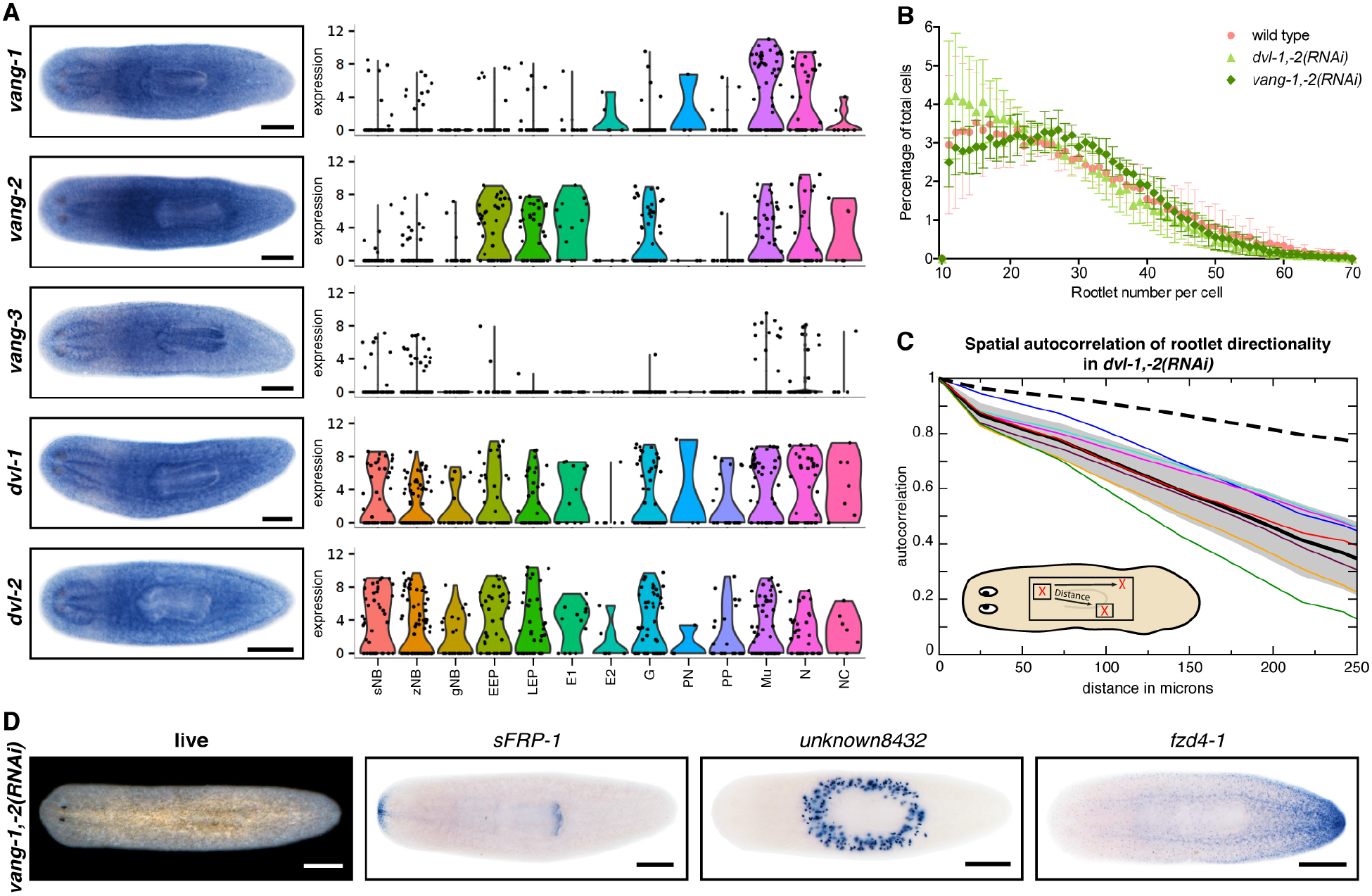
Core PCP is required for the A/P polarity component. **A.** Gene expression analysis of the planarian homologs of VanGogh (*vang*) and Dishevelled (*dvl*). Left panel: Whole mount gene expression patterns of the indicated genes by in situ hybridization. Scale bar: 500 μm. Right panel: Single cell gene expression profiles of the indicated genes in different cell types, generated using the Planarian SCS online resource (Wurtzel et al., 2015). sNB: sigma neoblast; zNB: zeta neoblast; gNB: gamma neoblast; EEP: early epidermal progenitors; LEP: late epidermal progenitors; E1: epidermis I; E2: epidermis 2; Gut: gut; PN: protonephridia; PP: parapharyngeal; Mu: muscle; N: neural; N-C: neural (ciliated). **B.** Lack of ciliogenesis phenotypes under PCP components knockdown. Histogram representation of rootlet number/epidermal cell of the same specimens that used for polarity analysis under *vang-1,-2(RNAi)*, *dvl-1,-2(RNAi),* and wild-type conditions. Rootlet number was identified using customized image analysis algorithm; epidermal cell boundary was determined using Voronoi tessellation (see Materials and Methods for detail explanation). **C.** Spatial autocorrelation analysis of rootlet directionality in 7 intact *dvl-1,-2(RNAi)* specimens, comparing directional coherence within square of 150×150 pixels. **D.** Body plan polarity landmarks in representative intact *vang-1,-2(RNAi)* specimens. Live image (top left), whole mount in situ hybridization gene expression patterns of head, trunk and tail markers. Scale bar: 500 μm. Normal marker expression indicates lack of overt body plan patterning phenotypes. Scale bar: 500 μm.

**Fig. S7:**
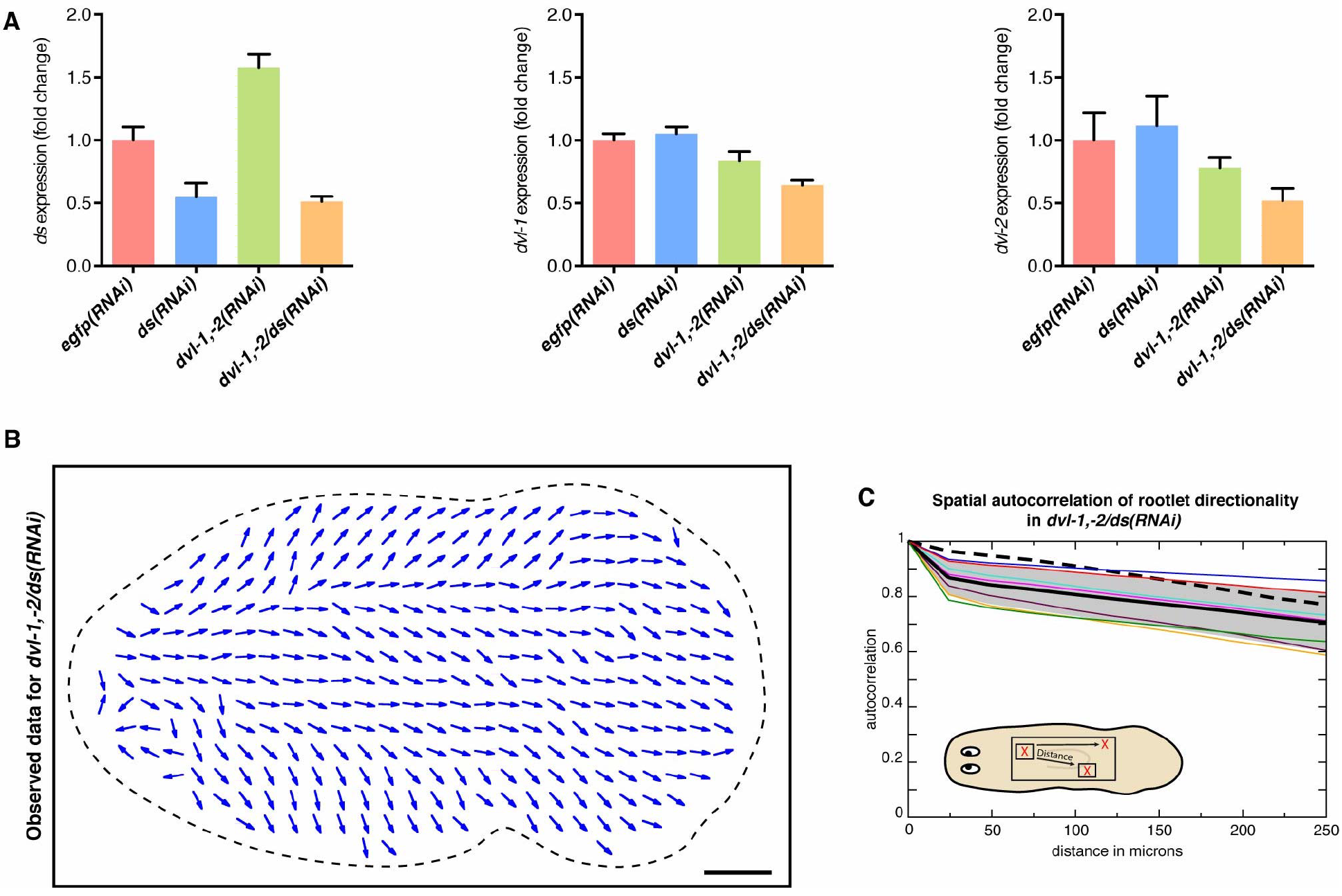
Ciliary polarity in the absence of both M/L and A/P components. **A.** Gene expression levels of *ds* (left), *dvl-1* (middle) and *dvl-2* (right) by qPCR under the indicated RNAi conditions. **B.** Vector field representation of the average ciliary rootlet orientation within 100 μm bins of a representative *dvl-1,-2/ds* triple-RNAi specimen. **C.** Spatial autocorrelation analysis of rootlet directionality in 7 intact *dvl-1,-2/ds(RNAi)* specimens, comparing directional coherence within square of 150×150 pixels.

## References

Adell, T., Saló, E., Boutros, M., and Bartscherer, K. (2009). Smed-Evi/Wntless is required for beta-catenin-dependent and -independent processes during planarian regeneration. Development 136, 905–910.

Adler, P.N. (2012). The frizzled/stan Pathway and Planar Cell Polarity in the Drosophila Wing. Curr. Top. Dev. Biol. 101, 1–31.

Aigouy, B., Farhadifar, R., Staple, D.B., Sagner, A., Röper, J.-C., Jülicher, F., and Eaton, S. (2010). Cell Flow Reorients the Axis of Planar Polarity in the Wing Epithelium of Drosophila. Cell 142, 773–786.

Alexandre, C., Baena-Lopez, A., and Vincent, J.-P. (2014). Patterning and growth control by membrane-tethered Wingless. Nature 505, 180–.

Almuedo-Castillo, M., Saló, E., and Adell, T. (2011). Dishevelled is essential for neural connectivity and planar cell polarity in planarians. Pnas 108, 2813–2818.

Alvarado, A.S. (2002). The Schmidtea mediterranea database as a molecular resource for studying platyhelminthes, stem cells and regeneration. Development 129, 5659–5665.

Amonlirdviman, K. (2005). Mathematical Modeling of Planar Cell Polarity to Understand Domineering Nonautonomy. Science 307, 423–426.

Aslanidis, C., and de Jong, P.J. (1990). Ligation-independent cloning of PCR products (LIC-PCR). Nucleic Acids Res. 18, 6069–6074.

Aw, W.Y., and Devenport, D. (2017). Planar cell polarity: global inputs establishing cellular asymmetry. Current Opinion in Cell Biology 44, 110–116.

Azimzadeh, J., Wong, M.L., Downhour, D.M., Alvarado, A.S., and Marshall, W.F. (2012). Centrosome Loss in the Evolution of Planarians. Science 335, 461–463.

Blassberg, R.A., Felix, D.A., Tejada-Romero, B., and Aboobaker, A.A. (2013). PBX/extradenticle is required to re-establish axial structures and polarity during planarian regeneration. Development 140, 730–739.

Blasse, C., Saalfeld, S., Etournay, R., Sagner, A., Eaton, S., and Myers, E.W. (2017). PreMosa: extracting 2D surfaces from 3D microscopy mosaics. Bioinformatics 33, 2563–2569.

Brittle, A., Thomas, C., and Strutt, D. (2012). Planar Polarity Specification through Asymmetric Subcellular Localization of Fat and Dachsous. Current Biology 22, 907–914.

Burak, Y., and Shraiman, B.I. (2009). Order and Stochastic Dynamics in Drosophila Planar Cell Polarity. PLoS Comput Biol 5.

Butler, M.T., and Wallingford, J.B. (2017). Planar cell polarity in development and disease. Nature Reviews Molecular Cell Biology 18, 375–388.

Cebrià, F., and Newmark, P.A. (2005). Planarian homologs of netrin and netrin receptor are required for proper regeneration of the central nervous system and the maintenance of nervous system architecture. Development 132, 3691–3703.

Chaikin, P.M., and Lubensky, T.C. (2000). Principles of Condensed Matter Physics (Cambridge University Press).

Chen, C.C.G., Wang, I.E., and Reddien, P.W. (2013). pbx is required for pole and eye regeneration in planarians. Development 140, 719–729.

Chien, Y.-H., Keller, R., Kintner, C., and Shook, D.R. (2015). Mechanical Strain Determines the Axis of Planar Polarity in Ciliated Epithelia. Current Biology 25, 2774–2784.

Chu, C.-W., and Sokol, S.Y. (2016). Wnt proteins can direct planar cell polarity in vertebrate ectoderm. Elife 5.

Clark, H.F., Brentrup, D., Schneitz, K., Bieber, A., Goodman, C., and Noll, M. (1995). Dachsous encodes a member of the cadherin superfamily that controls imaginal disc morphogenesis in Drosophila. Genes Dev. 9, 1530–1542.

Devenport, D., Oristian, D., Heller, E., and Fuchs, E. (2011). Mitotic internalization of planar cell polarity proteins preserves tissue polarity. Nat. Cell Biol. 13, 893–902.

Frisch, D., and Farbman, A.I. (1968). Development of order during ciliogenesis. Anat. Rec. 162, 221–232.

Gaviño, M.A., and Reddien, P.W. (2011). A Bmp/Admp Regulatory Circuit Controls Maintenance and Regeneration of Dorsal-Ventral Polarity in Planarians. Current Biology 21, 294–299.

Glazer, A.M., Wilkinson, A.W., Backer, C.B., Lapan, S.W., Gutzman, J.H., Cheeseman, I.M., and Reddien, P.W. (2010). The Zn Finger protein Iguana impacts Hedgehog signaling by promoting ciliogenesis. Dev. Biol. 337, 148–156.

Goodrich, L.V., and Strutt, D. (2011). Principles of planar polarity in animal development. Development 138, 1877–1892.

Gurley, K.A., Elliott, S.A., Simakov, O., Schmidt, H.A., Holstein, T.W., and Alvarado, A.S. (2010). Expression of secreted Wnt pathway components reveals unexpected complexity of the planarian amputation response. Dev. Biol. 347, 24–39.

Gurley, K.A., Rink, J.C., and Alvarado, A.S. (2008). β-catenin defines head versus tail identity during planarian regeneration and homeostasis. Science 319, 323–327.

Hale, R., and Strutt, D. (2015). Conservation of Planar Polarity Pathway Function Across the Animal Kingdom. Annu. Rev. Genet. 49, 529–551.

Hazelwood, L.D., and Hancock, J.M. (2013). Functional modelling of planar cell polarity: an approach for identifying molecular function. BMC Dev. Biol. 13.

Hoffmann, K.B., Voss-Boehme, A., Rink, J.C., and Brusch, L. (2017). A dynamically diluted alignment model reveals the impact of cell turnover on the plasticity of tissue polarity patterns. Journal of the Royal Society Interface 14.

Iglesias, M., Gomez-Skarmeta, J.L., Saló, E., and Adell, T. (2008). Silencing of Smed-catenin1 generates radial-like hypercephalized planarians. Development 135, 1215–1221.

King, R.S., and Newmark, P.A. (2013). In situ hybridization protocol for enhanced detection of gene expression in the planarian Schmidtea mediterranea. BMC Dev. Biol. 13, 8.

Lander, R., and Petersen, C.P. (2016). Wnt, Ptk7, and FGFRL expression gradients control trunk positional identity in planarian regeneration. Elife 5, e02238.

Larsson, M., Gräslund, S., Yuan, L., Brundell, E., Uhlén, M., Höög, C., and Ståhl, S. (2000). High-throughput protein expression of cDNA products as a tool in functional genomics. J. Biotechnol. 80, 143–157.

Lawrence, P.A., Struhl, G., and Casal, J. (2007). Planar cell polarity: one or two pathways? Nat. Rev. Genet. 8, 555–563.

Liu, S.Y., Selck, C., Friedrich, B., Lutz, R., Vila-Farre, M., Dahl, A., Brandl, H., Lakshmanaperumal, N., Henry, I., and Rink, J.C. (2013). Reactivating head regrowth in a regeneration-deficient planarian species. Nature 500, 81–84.

Ma, D., Yang, C.H., McNeill, H., Simon, M.A., and Axelrod, J.D. (2003). Fidelity in planar cell polarity signalling. Nature 421, 543–547.

Mao, Y.P., Rauskolb, C., Cho, E., Hu, W.L., Hayter, H., Minihan, G., Katz, F.N., and Irvine, K.D. (2006). Dachs: an unconventional myosin that functions downstream of Fat to regulate growth, affinity and gene expression in Drosophila. Development 133, 2539–2551.

Matakatsu, H. (2004). Interactions between Fat and Dachsous and the regulation of planar cell polarity in the Drosophila wing. Development 131, 3785–3794.

Matakatsu, H., and Blair, S.S. (2006). Separating the adhesive and signaling functions of the Fat and Dachsous protocadherins. Development 133, 2315–2324.

Matakatsu, H., and Blair, S.S. (2012). Separating planar cell polarity and Hippo pathway activities of the protocadherins Fat and Dachsous. Development 139, 1498–1508.

Matis, M., and Axelrod, J.D. (2013). Regulation of PCP by the Fat signaling pathway. Genes Dev. 27, 2207–2220.

Matis, M., Russler-Germain, D.A., Hu, Q., Tomlin, C.J., and Axelrod, J.D. (2014). Microtubules provide directional information for core PCP function. Elife 3, –17.

Oderberg, I.M., Li, D.J., Scimone, M.L., Gaviño, M.A., and Reddien, P.W. (2017). Landmarks in Existing Tissue at Wounds Are Utilized to Generate Pattern in Regenerating Tissue. Current Biology 27, 733–742.

Otsu, N. (1979). A Threshold Selection Method from Gray-Level Histograms. IEEE Transactions on Systems, Man, and Cybernetics 9, 62–66.

Park, T.J., Mitchell, B.J., Abitua, P.B., Kintner, C., and Wallingford, J.B. (2008). Dishevelled controls apical docking and planar polarization of basal bodies in ciliated epithelial cells. Nat Genet 40, 871–879.

Pearson, B.J., Eisenhoffer, G.T., Gurley, K.A., Rink, J.C., Miller, D.E., and Sánchez Alvarado, A. (2009). Formaldehyde-based whole-mount in situ hybridization method for planarians. Dev. Dyn. 238, 443–450.

Petersen, C.P., and Reddien, P.W. (2008). Smed- catenin-1 Is Required for Anteroposterior Blastema Polarity in Planarian Regeneration. Science 319, 327–330.

Petersen, C.P., and Reddien, P.W. (2009). A wound-induced Wnt expression program controls planarian regeneration polarity. Pnas 106, 17061–17066.

Petersen, C.P., and Reddien, P.W. (2011). Polarized notum Activation at Wounds Inhibits Wnt Function to Promote Planarian Head Regeneration. Science 332, 852–855.

Reuter, H., März, M., Vogg, M.C., Eccles, D., Grífol-Boldú, L., Wehner, D., Owlarn, S., Adell, T., Weidinger, G., and Bartscherer, K. (2015). β-Catenin-Dependent Control of Positional Information along the AP Body Axis in Planarians Involves a Teashirt Family Member. Cell Rep 10, 253–265.

Rieger, R.M. (1981). Morphology of the Turbellaria at the Ultrastructural Level. Hydrobiologia 84, 213–229.

Rink, J.C., Gurley, K.A., Elliott, S.A., and Sánchez Alvarado, A. (2009). Planarian Hh Signaling Regulates Regeneration Polarity and Links Hh Pathway Evolution to Cilia. Science 326, 1406–1410.

Rink, J.C. (2012). Stem cell systems and regeneration in planaria. Dev. Genes Evol. 223, 67–84.

Rompolas, P., Azimzadeh, J., Marshall, W.F., and King, S.M. (2013). Analysis of Ciliary Assembly and Function in Planaria. Meth. Enzymol. 525, 245–264.

Rouhana, L., Weiss, J.A., Forsthoefel, D.J., Lee, H., King, R.S., Inoue, T., Shibata, N., Agata, K., and Newmark, P.A. (2013). RNA interference by feeding in vitro-synthesized double-stranded RNA to planarians: Methodology and dynamics. Dev. Dyn. 242, 718–730.

Sagner, A., Merkel, M., Aigouy, B., Gaebel, J., Brankatschk, M., Juelicher, F., and Eaton, S. (2012). Establishment of Global Patterns of Planar Polarity during Growth of the Drosophila Wing Epithelium. Current Biology 22, 1296–1301.

Sánchez Alvarado, A., and Newmark, P.A. (1999). Double-stranded RNA specifically disrupts gene expression during planarian regeneration. Pnas 96, 5049–5054.

Schindelin, J., Arganda-Carreras, I., Frise, E., Kaynig, V., Longair, M., Pietzsch, T., Preibisch, S., Rueden, C., Saalfeld, S., Schmid, B., et al. (2012). Fiji: an open-source platform for biological-image analysis. Nat. Methods 9, 676–682.

Scimone, M.L., Cote, L.E., Rogers, T., and Reddien, P.W. (2016). Two FGFRL-Wnt circuits organize the planarian anteroposterior axis. Elife 5, 905.

Smith, C.S., Preibisch, S., Joseph, A., Abrahamsson, S., Rieger, B., Myers, E., Singer, R.H., and Grunwald, D. (2015). Nuclear accessibility of β-actin mRNA is measured by 3D single-molecule real-time tracking. J. Cell Biol. 209, 609–619.

Sonka, M., Hlavac, V., and Boyle, R. (2007). Image Processing, Analysis and Machine Vision 3rd Edition (Thomson Engineering).

Strutt, D. (2009). Gradients and the Specification of Planar Polarity in the Insect Cuticle. Cold Spring Harb Perspect Biol 1, –a000489.

Stückemann, T., Cleland, J.P., Werner, S., Vu, H.T.-K., Bayersdorf, R., Liu, S.-Y., Friedrich, B., Jülicher, F., and Rink, J.C. (2017). Antagonistic Self-Organizing Patterning Systems Control Maintenance and Regeneration of the Anteroposterior Axis in Planarians. Dev. Cell 40, 248–263.e4.

Sureda-Gómez, M., Martín-Durán, J.M., and Adell, T. (2016). Localization of planarian β-CATENIN-1 reveals multiple roles during anterior-posterior regeneration and organogenesis. Development 143, 4149–4160.

Szeliski, R. (2011). Computer Vision (London: Springer London).

Taylor, J., Abramova, N., Charlton, J., and Adler, P.N. (1998). Van Gogh: A new Drosophila tissue polarity gene. Genetics 150, 199–210.

Tu, K.C., Cheng, L.-C., TK Vu, H., Lange, J.J., McKinney, S.A., Seidel, C.W., and Sánchez Alvarado, A. (2015). Egr-5 is a post-mitotic regulator of planarian epidermal differentiation. Elife 4, e02238.

van Wolfswinkel, J.C., Wagner, D.E., and Reddien, P.W. (2014). Single-Cell Analysis Reveals Functionally Distinct Classes within the Planarian Stem Cell Compartment. Cell Stem Cell 15, 326–339.

Vásquez-Doorman, C., and Petersen, C.P. (2014). zic-1 Expression in Planarian Neoblasts after Injury Controls Anterior Pole Regeneration. PLoS Genet. 10, e1004452.

Villano, J.L., and Katz, F.N. (1995). four-jointed is required for intermediate growth in the proximal-distal axis in Drosophila. Development 121, 2767–2777.

Vinson, C.R., and Adler, P.N. (1987). Directional non-cell autonomy and the transmission of polarity information by the frizzled gene of Drosophila. Nature 329, 549–551.

Vogg, M.C., Owlarn, S., Pérez Rico, Y.A., Xie, J., Suzuki, Y., Gentile, L., Wu, W., and Bartscherer, K. (2014). Stem cell-dependent formation of a functional anterior regeneration pole in planarians requires Zic and Forkhead transcription factors. Dev. Biol. 390, 136–148.

Wallingford, J.B. (2010). Planar cell polarity signaling, cilia and polarized ciliary beating. Current Opinion in Cell Biology 22, 597–604.

Wu, J., Roman, A.-C., Carvajal Gonzalez, J.M., and Mlodzik, M. (2013). Wg and Wnt4 provide long-range directional input to planar cell polarity orientation in Drosophila. Nat. Cell Biol. 15, 1045–.

Wurtzel, O., Cote, L.E., Poirier, A., Satija, R., Regev, A., and Reddien, P.W. (2015). A Generic and Cell-Type-Specific Wound Response Precedes Regeneration in Planarians. Dev. Cell 35, 632–645.

Yamashita, S., and Michiue, T. (2016). Boundary propagation of planar cell polarity is robust against cell packing pattern. J. Theor. Biol. 410, 44–54.

Yang, J., Liu, X.Q., Yue, G.H., Adamian, M., Bulgakav, O., and Li, T.S. (2002). Rootletin, a novel coiled-coil protein, is a structural component of the ciliary rootlet. J. Cell Biol. 159, 431–440.

Yang, Y., and Mlodzik, M. (2015). Wnt-Frizzled/Planar Cell Polarity Signaling: Cellular Orientation by Facing the Wind (Wnt). Annu. Rev. Cell Dev. Biol. 31, 623–646.

Zeidler, M.P., Perrimon, N., and Strutt, D.I. (2000). Multiple roles for four-jointed in planar polarity and limb patterning. Dev. Biol. 228, 181–196.

## Supplemental material references

Eaton, S., and Jülicher, F. (2011). Cell flow and tissue polarity patterns. Curr. Opin. Genet. Dev. 21, 747–752.

Merkel, M., Sagner, A., Gruber, F.S., Etournay, R., Blasse, C., Myers, E., Eaton, S., and Juelicher, F. (2014). The Balance of Prickle/Spiny-Legs Isoforms Controls the Amount of Coupling between Core and Fat PCP Systems. Current Biology 24, 2111–2123.

Rappel, W.-J., and Edelstein-Keshet, L. (2017). Mechanisms of Cell Polarization. Curr Opin Syst Biol 3, 43–53.

Starruy, J., de Back, W., Brusch, L., and Deutsch, A. (2014). Morpheus: a user-friendly modeling environment for multiscale and multicellular systems biology. Bioinformatics 30, 1331–1332.

Szeliski, R. (2011). Computer Vision (London: Springer London).

